# Genetic Interplay Between Transcription Factor Pou4f1/Brn3a and Neurotrophin Receptor Ret In Retinal Ganglion Cell Type Specification

**DOI:** 10.1101/2020.03.23.004242

**Authors:** Vladimir Vladimirovich Muzyka, Tudor Constantin Badea

**Affiliations:** Retinal Circuit Development & Genetics Unit, Neurobiology-Neurodegeneration & Repair Laboratory, National Eye Institute, NIH, Bethesda, MD

**Author notes:** corresponding author: Tudor C. Badea / Vladimir V. Muzyka, Retinal Circuit Development & Genetics Unit, Building 6, Room 331, 6 Center Drive, Bethesda, MD 20892-0610, or /.

## Abstract

While the transcriptional code governing retinal ganglion cell (RGC) type specification begins to be understood, its interplay with neurotrophic signaling is largely unexplored. Using sparse random recombination, we show that mosaic gene dosage manipulation of the transcription factor Brn3a/Pou4f1 in neurotrophic receptor Ret heterozygote RGCs results in altered cell fate decisions and/or morphological dendritic defects. Specific RGC types are lost if Brn3a is ablated during embryogenesis and only mildly affected by postnatal Brn3a ablation. Sparse but not complete Brn3a heterozygosity combined with complete Ret heterozygosity has striking effects on RGC type distribution. Brn3a only mildly modulates Ret transcription, while Ret knockouts exhibit normal Brn3a and Brn3b expression. However, Brn3a loss of function significantly affects distribution of Ret co-receptors GFRα1-3, and neurotrophin receptors TrkA and TrkC in RGCs. Based on these observations, we propose that Brn3a and Ret converge onto developmental pathways that control RGC type specification, potentially through a competitive mechanism requiring signaling from the surrounding tissue.

## 1. Introduction

Retinal Ganglion Cells (RGCs) – the output neurons of the vertebrate retina – relay visual information to distinct projection areas in the brain. Currently, mouse RGCs are subdivided into about 50 types based on classification criteria including morphological, functional, and molecular parameters ^1–10^. The developmental mechanisms orchestrating the differentiation of RGC types involve transcription factors (TFs) in combination with extracellular signaling. Within the retina, Atoh7/Math5 ^11–13^ is required but not sufficient for neuronal precursors to commit to the RGC fate. Downstream of Atoh7, postmitotic TFs determine general traits of neuronal morphology and functional characteristics – in RGCs this group includes the three members of the Pou4f family, namely Pou4f1/Brn3a, Pou4f2/Brn3b, and Pou4f3/Brn3c ^2,14–20^. Pou4f/Brn3 TFs are expressed in the retina specifically in RGCs. Brn3a transcription is initiated somewhat later than Brn3b (embryonic day 13.5 – E13.5 vs embryonic day 11.5 – E11.5), acts downstream of Brn3b in the developmental transcriptional program, and was initially considered to function redundantly with Brn3b^21^.

Deletion of Brn3a in mice leads to early postnatal lethality caused by somatosensory system and brainstem nuclei abnormalities, with no gross perturbations in the retina at this stage of development^19,22^. Using a conditional alkaline phosphatase (AP) reporter allele knocked-in to the Brn3a locus (*Brn3a*^*CKOAP*^) we demonstrated that Brn3a-expressing RGCs laminate in the outer strata (~70%) of the inner plexiform layer (IPL) of the retina. Loss of Brn3a before the actual onset of locus expression results in a shift towards bistratified arbor morphologies and general decrease of RGC numbers by approximately 30% ^2,14,17^. This reduction is mostly explained by a loss of RGCs with small dendritic arbor areas and dense multistratified lamination pattern – ON and OFF β RGCs. RNA deep sequencing of affinity-purified early postnatal Brn3a^AP/KO^ (effectively Brn3a null = Brn3a^KO/KO^) RGCs revealed potential transcriptional targets for Brn3a regulation of cell type development ^9,23^. However, at what developmental stage Brn3a is necessary for specification of ON and OFF β RGCs still remains unexplored.

Amongst the neurotrophic cues required for neuronal development and specification, target derived neurotrophin (NT) and glial derived neurotrophic factor (GDNF) families of ligands play a major role. Components of receptor complexes for NTs contain members of distinct molecular families such as Trk, p75^NTR^ and sortilin ^24–27^. Neurotrophin receptors are also expressed in rodent RGCs during development ^9,28^ and development of dendrites and axons and physiological maturation of RGC is modulated by neurotrophin-3 (NT-3), brain derived neurotrophic factor (BDNF), and neurotrophin receptors, TrkB and p75^NTR 29–37^. Not much is known about the effects of Glial family ligands (GFLs) and their receptors in RGC development and specification. The GDNF family of neurotrophins contains four members – GDNF, artemin, neurturin, and persephin. Four co-receptors, “GDNF family receptor-α” (GFRα 1-4) attached to the cell membrane through a Glycosyl Phosphatidylinositol (GPI) anchor have selective affinity to the four ligands ^38–41^. The tyrosine kinase Ret co-receptor is required for downstream signaling through GFRα. Ret ablation phenotype is characterized by dramatic abnormalities of kidney formation, severely affected sympathetic ganglia, and defects in specific pain and touch somatosensory receptor cells, and megacolon (Hirschsprung disease) due to defects in the specification and migration of cells of the enteric nervous system ^42–47^. GFRα1, GFRα2 and Ret are expressed in RGC subpopulations, however, no RGC phenotypes were reported in Ret or GFR mutants. Ret and Neurturin mutants affect photoreceptor light responses and lead to morphological defects of photoreceptors, bipolar and horizontal cell contacts in the outer plexiform layer ^48^. Ret is dynamically expressed in specific retinal cell populations, beginning with RGCs (E13 – 14.5), followed by Horizontal (E17) and Amacrine (P1) cells ^49–51^. In the adult retina, Ret is co-expressed with Brn3a predominantly in three mono- and two bistratified subtypes of RGCs, with Brn3b – in four mono- and two bistratified subtypes, and with Brn3c – in a single monostratified subpopulation. Of interest, Brn3a and Ret are co-expressed in ON and OFF β RGCs ^51^.

In the current study, we use a *Ret*^*CreERt2*^ allele to induce sparse random recombination in Cre-dependent histochemical reporters targeted at the Brn3a and Rosa26 loci, to visualize RGC dendritic arbor morphologies in sparse or complete double heterozygote (*Ret*^*KO/WT*^; *Brn3a*^*KO/WT*^) retinas at different developmental stages. In addition, we reveal the effects of knocking out Brn3a at important developmental timepoints on RGC subtype distribution. Finally, using immunostaining in Ret or Brn3a complete retinal knock-outs, we assess the potential crosstalk between transcriptional and neurotrophic mechanisms in RGCs. We find that Brn3a is required for the development of at least two monostratified RGC subtypes during embryonic and perinatal stages. However, the most striking finding is the specific effect of embryonic sparse double-heterozygosity on cell type specification in mono- and bistratified RGCs and on dendritic morphology in a subset of bistratified RGCs.

## 2. Materials and Methods

### 2.1. Mouse lines and crosses

Mouse lines carrying *Rosa26*^*iAP* 52^, *Brn3a*^*KO* 19^, *Brn3a*^*CKOAP* 14^, *Ret*^*CreERt2* 44^, *Rax:Cre* ^53^ and *Ret*^*CKCFP* 47^ alleles were previously described. In the *Brn3a*^*KO*^ allele, the entire open reading frame of Brn3a is deleted and replaced with a Neo cassette. *Brn3a*^*KO/KO*^ mice are perinatal lethal, while *Brn3a*^*KO/WT*^ mice are viable and breed normally ^19^. In the *Brn3a*^*CKOAP*^ allele the two coding exons of Brn3a are appended with a (3x SV40) transcriptional STOP and flanked by loxP sites ^14^ and the cDNA of the histochemical reporter Alkaline Phosphatase (AP) is inserted after the 3’ loxP site. Cre mediated recombination results in ablation of the Brn3a ORF coupled with expression of the AP cDNA under the control of the endogenous regulatory elements of the Brn3a locus. The *Rosa26*^*iAP*^ reporter locus expresses AP ubiquitously in a Cre dependent manner ^52^. The *Ret*^*CKCFP*^ conditional minigene allele includes the complete human Ret cDNA flanked by loxP sites knocked-in into exon 1 of the mouse *Ret* gene and followed by the Cyan Fluorescent Protein (CFP) cDNA^47^. The unrecombined locus expresses the full human Ret ORF, while Cre mediated recombination ablates Ret, and replaces it with CFP, generating a Ret null allele. The BAC transgenic mouse line Rax:Cre contains Cre recombinase controlled by the mouse *Rax* gene locus ^53^, and expresses Cre in the anterior eye field, beginning with E9. The *Ret*^*CreERt2*^ allele contains the CreERt2 (tamoxifen-dependent Cre activity) coding sequence knocked-in in the first exon of the *Ret* gene ^44^, resulting in a Ret null allele.

To understand the cell-autonomous effects of losing one or both copies of Brn3a at different time points from Ret^+^ RGCs, we have crossed *Ret*^*CreERt2/WT*^; *Brn3a*^*KO/WT*^ males x *Brn3a*^*CKOAP/CKOAP*^ females resulting in *Ret*^*CreERt2/WT*^; *Brn3a*^*CKOAP/WT*^ and *Ret*^*CreERt2/WT*^; *Brn3a*^*CKOAP/KO*^ pups. To achieve random sparse recombination and AP expression, we induced Cre recombinase by intraperitoneal (i.p.) injection of 4-hydroxytamoxifen (4-HT) at postnatal day 22 (P22 – adult, 50-100 μg 4-HT), postnatal day 0 (P0 pups, 2.5-5 μg 4-HT), or at embryonic day 15 (E15 embryos, delivered i.p. to the mother, 250 μg 4-HT) (Figure 1 a, b). To study the effects of complete double-heterozygosity (Brn3a^KO/WT^; Ret^KO/WT^) on RGC dendritic morphology, we crossed *Ret*^*CreERt2/WT*^; *Brn3a*^*KO/WT*^ males x *ROSA26*^*AP/AP*^ females to get *Ret*^*CreERt2/WT*^; *Brn3a*^*KO/WT*^; *ROSA26*^*AP/WT*^ and *Ret*^*CreERt2/WT*^; *Brn3a*^*WT/WT*^; *ROSA26*^*AP/WT*^ pups. Cre recombination and AP expression were induced by i.p. injections of 4-HT to the mother at E15 (250 μg 4-HT, Figure 1 c, d). For each genotype, and age of 4HT treatment, retinas from at least three animals were analyzed at two months after injection, with the exception of P0 induced *Ret*^*CreERt2/WT*^; *Brn3a*^*CKOAP/WT*^ mice, where only two animals were analyzed.

**Figure 1.**
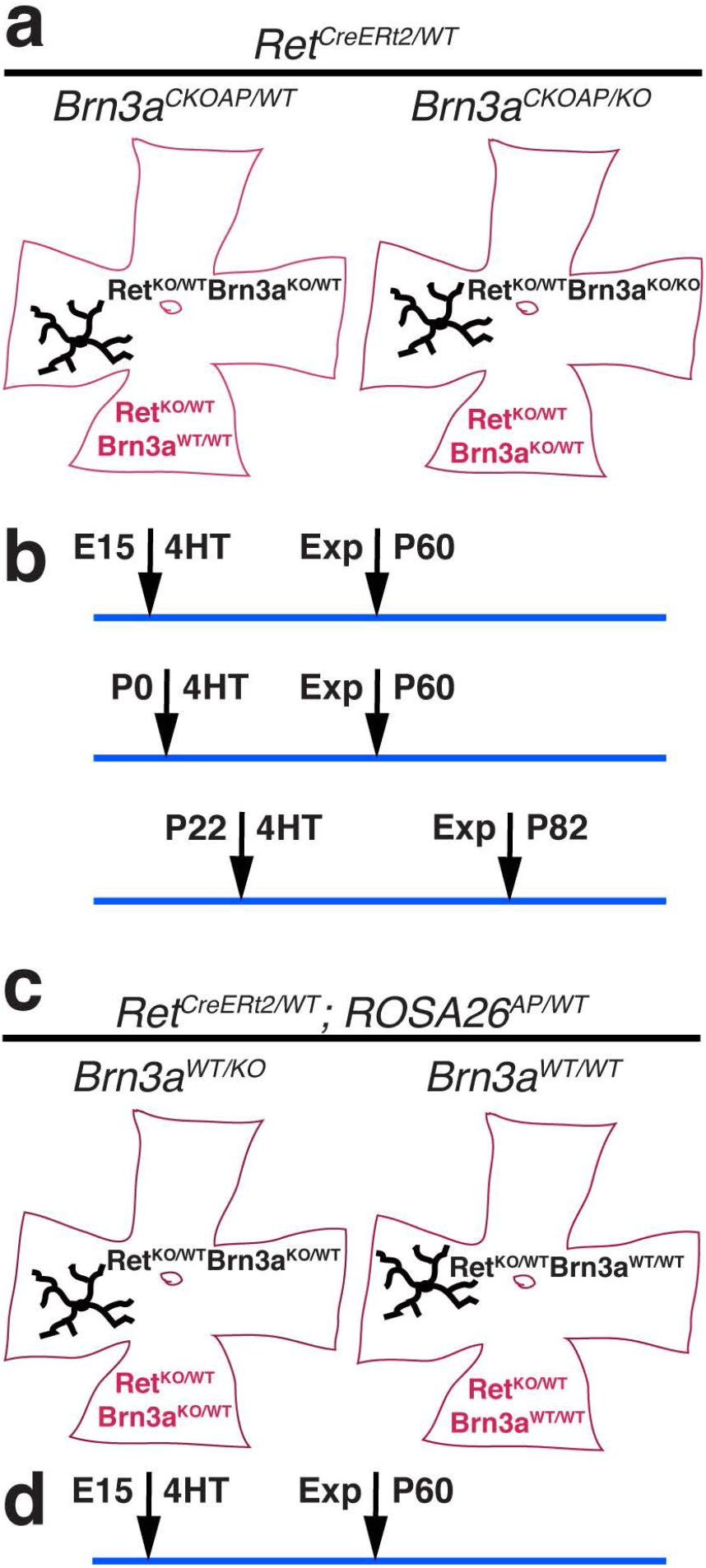
Mouse genotypes and timelines for experiments in figures 2 through 6. a, Retinas from *Ret*^*CreERt2/WT*^; *Brn3a*^*CKOAP/WT*^ (left) and *Ret*^*CreERt2/WT*^; *Brn3a*^*CKOAP/KO*^ (right) mice, were injected with 4-HT to induce sparse random recombination. Recombined cells (black arbors) are either Ret^KO/WT^; Brn3a^KO/WT^ (Ret^CreERt2/WT^; Brn3a^AP/WT^) on a Ret^KO/WT^; Brn3a^WT/WT^ background (left) or Ret^KO/WT^; Brn3a^KO/KO^ (Ret^CreERt2/WT^; Brn3a^AP/KO^) on a Ret^KO/WT^; Brn3a^KO/WT^ background (right). b, Timelines for experiments in a. Injections were i.p. either to the pregnant female (for E15), to postnatal pups (P0) or adult mice (P22). Retinas from E15 or P0 inductions were stained at two months postnatal (P60), while P22 induced animals were analyzed at 2 months post injection (P82). c, Sparse random recombination approach in *Ret*^*CreERt2/WT*^; *Rosa26*^*iAP/WT*^; *Brn3a*^*KO/WT*^ (left) and *Ret*^*CreERt2/WT*^; *Rosa26*^*iAP/WT*^; *Brn3a*^*WT/WT*^ (right) retinas. After 4-HT induced recombination, RGCs are labelled by the *Rosa26*^*iAP/WT*^ allele, but preserve the same genotype as the surrounding tissue: Ret^KO/WT^; Brn3a^KO/WT^ (left) or Ret^KO/WT^; Brn3a^WT/WT^ (right). d, Timelines for experiments in c. Injections at E15 were i.p. to the pregnant female, and retinas of pups were stained at two months postnatal (P60).

**Figure 2.**
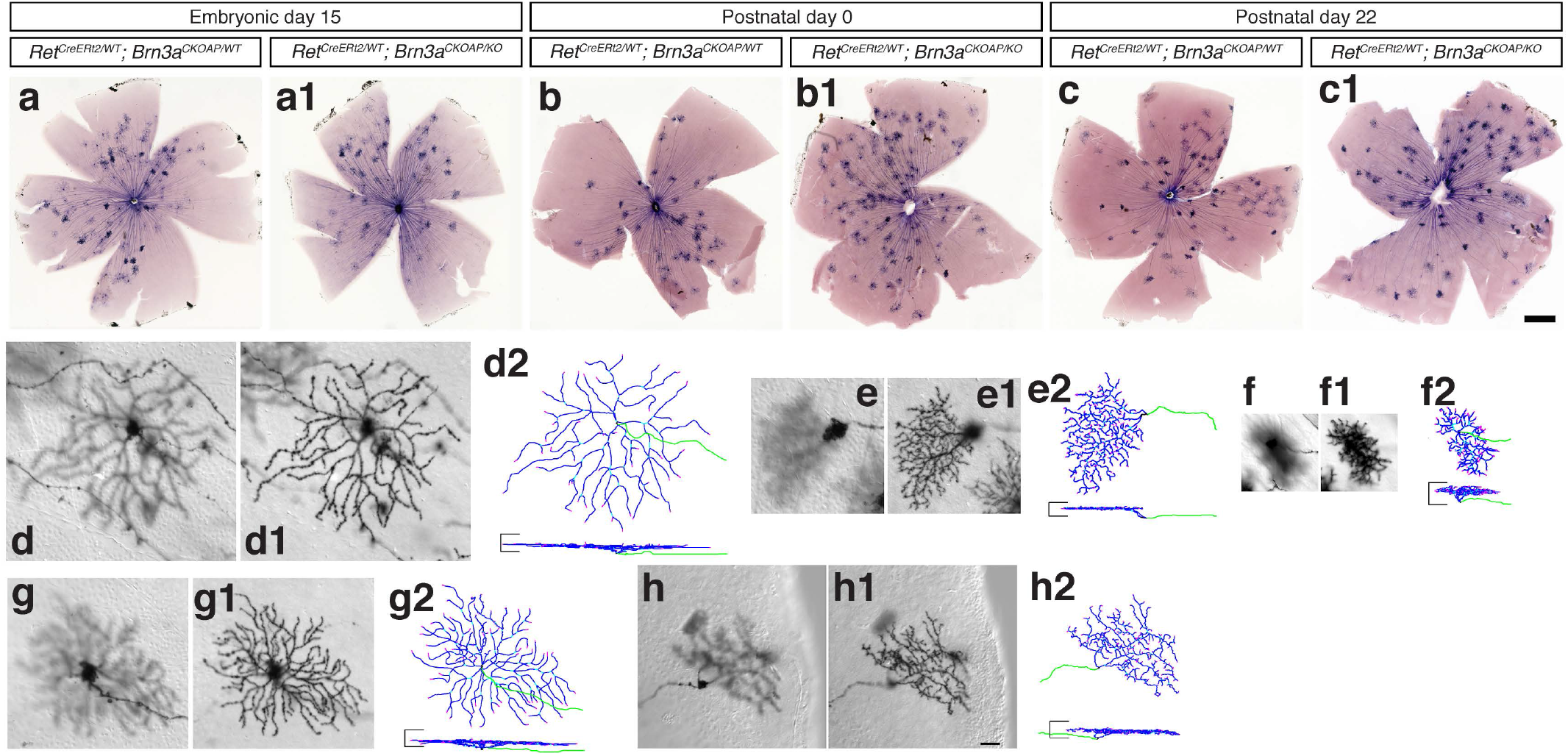
Monostratified RGCs in Ret^CreERt2^ Brn3a^CKOAP^ mouse retinas. (a-c1) AP histochemical staining of retinal flat mounts from *Ret*^*CreERt2/WT*^; *Brn3a*^*CKOAP/WT*^ (a, b, c) and *Ret*^*CreERt2/WT*^; *Brn3a*^*CKOAP/KO*^ (a1, b1, c1) mice, injected i.p. with 4-hydroxytamoxifen (4-HT) at embryonic day 15 (E15, mother injected i.p. at 15^th^ day of gestation with 250 μg 4-HT, a-a1), at postnatal day 0 (P0, 0.005 μg 4-HT, b-b1), or at postnatal day 22 (P22, 75-100 μg 4-HT, c-c1). (d-h2) Examples of most frequently encountered monostratified RGC morphologies: ONαS (d-d2), ON spiny (e-e2), ON and OFF betta (f-f2), M5 (g-g2) and Brn3aKO-specific betta (h-h2). For each cell, focal planes at the level of the cell body and axon (d, e, f, g, h), and dendritic arbor (d1, e1, f1, g1, h1) are shown in flat mount (en face) perspective. (d2, e2, f2, g2, h2) represent 3D reconstructions with en face perspective (top) and lateral projection (bottom). Axons are green, dendrites are blue. The horizontal bars on the sides of each vertical projection represent the boundaries of the IPL and are 25 μm long. Scale bar in (a-c1) is 500 μm, in h1 – 25 μm.

To study potential transcriptional regulation of Brn3a via Ret signaling we crossed Rax:Cre; *Ret*^*CKCFP/WT*^ males x *Ret*^*CKCFP/CKCFP*^ females resulting in Rax:Cre; *Ret*^*CKCFP/WT*^ (full-retina Ret-heterozygote) and Rax:Cre; *Ret*^*CKCFP/CKCFP*^ (full-retina Ret-knockout) offspring (Figure 7). To study potential Brn3a transcriptional regulation of Ret, GFRα and Trk receptors, we crossed Rax:Cre; *Brn3a*^*KO/WT*^ males x *Brn3a*^*CKOAP/CKOAP*^ females to get Rax:Cre; *Brn3a*^*CKOAP/WT*^ (full-retina Brn3a-heterozygote) and Rax:Cre; *Brn3a*^*CKOAP/KO*^ (full-retina Brn3a-knockout) offspring (Figure 7–9). For these experiments, tissues were harvested from mice of both sexes, between two and four months of age.

All mice were on C57/Bl6-SV129 mixed background. All animal procedures were approved by the National Eye Institute (NEI) Animal Care and Use Committee under protocol NEI640.

### 2.2. AP histochemistry and morphometric analysis

Mouse retinas were stained, processed, and imaged, and RGC dendritic arbors were traced and quantified as described previously ^1,51,54^. Animals were anesthetized and fixed by intracardiac perfusion with 4% Paraformaldehyde (PFA). Retinas were dissected and flat mounted, heat inactivated in a water bath at 65°C for one hour, and then AP histochemical stain developed. Color images of retina whole mounts and DIC grayscale image stacks (at 1 μm z step) of individual RGC dendritic arbors were captured with a Zeiss Imager.Z2. Morphological characteristics were measured using ImageJ software as described in Figure 3 and references ^1,51^. Relative lamination levels of dendritic arbors in the IPL were described by the lamination measurements in Figures 3, 5 and 6, and oriented by the previously reported stratification levels of ON and OFF Starburst Amacrine Cells (SACs) and the borderline between ON and OFF sublaminae of the IPL, as inferred from the lamination of axon terminals of ON bipolar cells ^55^. Neuronal reconstructions were made using Neuromantic (Darren Myat, http://www.reading.ac.uk/neuromantic) and projections were generated using a Matlab (Mathworks, Inc.) script ^17^. For each genotype combination and condition, at least three mice were analyzed.

**Figure 3.**
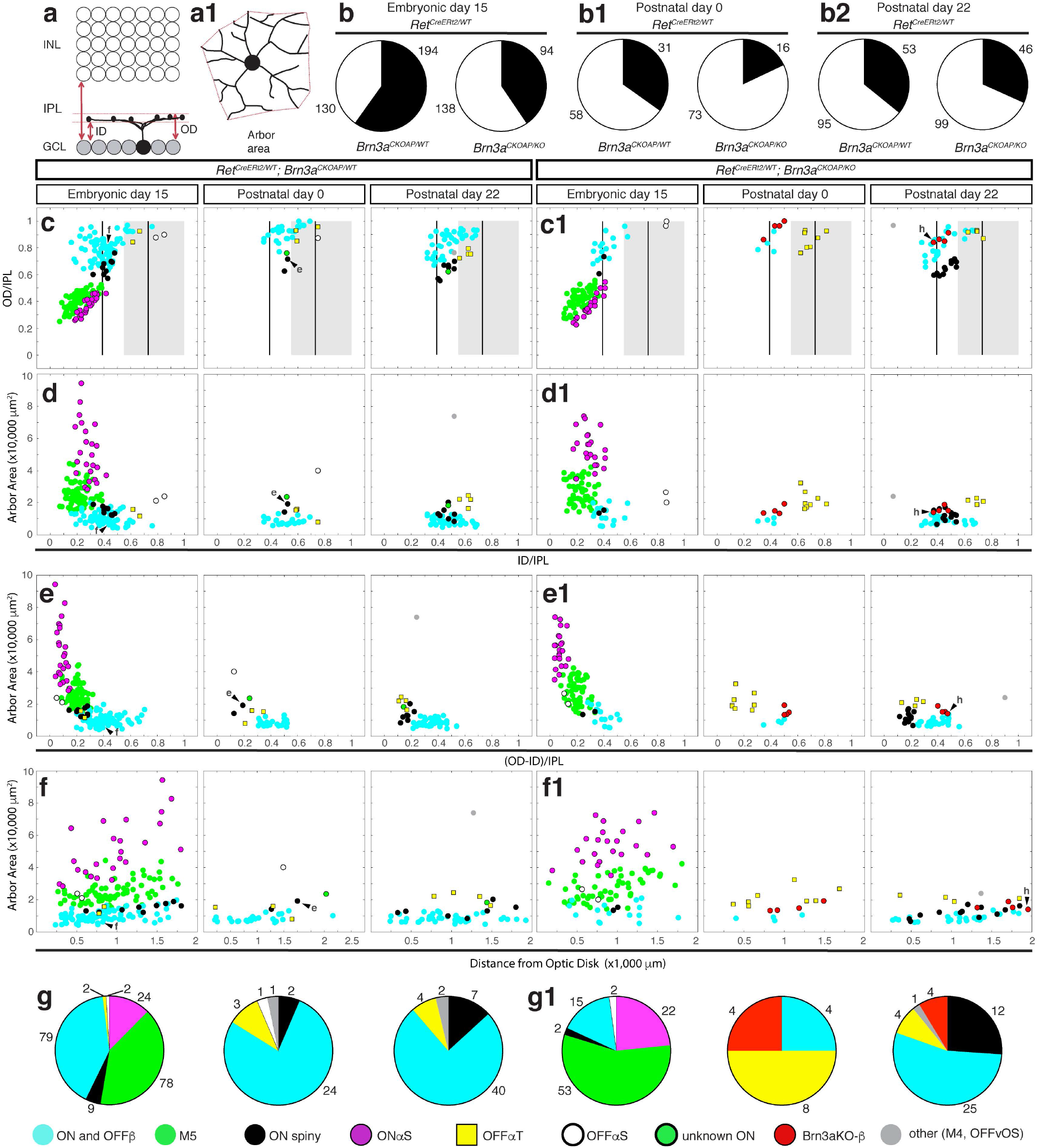
Morphological paramaters of Ret^CreERt2/WT^; Brn3a^AP/WT^ and Ret^CreERt2/WT^; Brn3a^AP/KO^ monostratified RGCs. (a-a1) RGC dendritic arbor parameters. (a) Inner distance (ID) and outer distance (OD), normalized to the IPL thickness, define the boundaries of dendritic arbor stratification, and are set to 0 at the GCL and 1 at the INL. (a1) Arbor area of the bounding polygon is determined from en face perspective. (b-b2) Pie-charts for proportions of monostratified (black) to bistratified (white) RGCs in *Ret*^*CreERt2/WT*^; *Brn3a*^*CKOAP/WT*^ (left) and *Ret*^*CreERt2/WT*^; *Brn3a*^*CKOAP/KO*^ (right) retinas of animals i.p. injected with 4-HT at E15 (b), P0 (b1), or P22 (b2). Numbers represent the total number of quantified RGCs for each genotype. (c-f1) Scatter plots of morphological parameters for monostratified RGCs. Ret^CreERt2/WT^; Brn3a^AP/WT^ cells – c, d, e, f; Ret^CreERt2/WT^; Brn3a^AP/KO^ cells – c1, d1, e1, f1. (c-c1) Normalized ID (ID/IPL, x) versus OD (OD/IPL, y). (d-d1) Normalized ID (ID/IPL, x) versus area (y). (e-e1) Normalized dendritic arbor thickness ((OD – ID)/IPL, x axis) versus arbor area (y). (f-f1) Distance from optic disc to the cell body (x axis) versus area (y). The two vertical bars in (c-c1) represent the theoretical lamination levels of ON and OFF starburst ACs, the grey box represents the OFF sublaminae of the IPL, where mGluR6^GFP^ positive axons are absent (Morgan et. al, 2006). The putative morphological types are assigned based on previous work and correspondence to literature, color coded in the scatter plots, and indicated at the bottom. Example cells from Figure 2 are indicated by arrowheads and corresponding letters. (g-g1) Pie-charts for proportions of different monostratified RGC subtypes in *Ret*^*CreERt2/WT*^; *Brn3a*^*CKOAP/WT*^ (g) and *Ret*^*CreERt2/WT*^; *Brn3a*^*CKOAP/KO*^ (g1) retinas. Numbers in (g-g1) represent the total number of quantified RGCs for each genotype and RGC type.

### 2.3. Indirect immunofluorescence

Retina vertical sections were processed and immunostained as previously described ^14,17,51^. In brief, retinas were fixed for 30 min in 2% paraformaldehyde, cryoprotected in OCT, and sectioned at 14 μm thickness on a cryostat. For each genotype, retinas from at least three different animals were sectioned and stained together on the same slide. Images (40x) were acquired using a Zeiss LSM700 confocal microscope and Zen software. Number of analyzed retinas and collected images are indicated in legend of Figure 7. Antibodies and dilutions used for analysis: 1:50 rabbit polyclonal anti-Brn3b generated in our lab ^51^; 1:20 mouse monoclonal anti-Brn3a (Millipore, MAB1585, RRID: AB_94166, clone 5A3.2; ^20^; 1:25 rabbit polyclonal anti-Ret (IBL, cat # R787); 1:50 mouse monoclonal anti-alkaline phosphatase (VEB Gent, Belgium, E6 clone); 1:40 goat polyclonal anti-TrkA (R&D Systems, AF1056); 1:40 goat polyclonal anti-TrkB (R&D Systems, AF1494); 1:40 goat polyclonal anti-TrkC (R&D Systems, AF1404); 1:20 goat polyclonal anti-GFRα1 (R&D Systems, AF560-SP); 1:100 goat polyclonal anti-GFRα2 (R&D Systems, AF429); 1:40 goat polyclonal anti-GFRα3 (R&D Systems, AF2645); 1:1000 chicken anti-GFP (used for detection of CFP protein; Abcam, ab13970, RRID: AB_300798). Alexa-Fluor conjugated Donkey polyclonal secondary antibodies were from Molecular Probes/Life Sciences and used at 1:300 dilution.

### 2.4. Statistical Methods

For RGC type distributions (Figures 2–6), data was collected from retinas from multiple animals for each treatment and genotype, and total numbers of measured cells are indicated in pie chart summaries in Figures 3, 5, and 6 and ranged from 90 to more than 300. Differences in cell type distribution were assessed using the Chi-square method, and Chi Statistics and P values indicated in supplementary table 1. For Indirect Immunofluorescence Experiments (Figures 7–9), data was collected from at least three animals, and cells were counted in 7 – 20 images for each genotype. Total numbers of measured cells are indicated in pie charts. Individual comparisons between groups of interest were performed using the Kolmogorov-Smirnov (KS2) test, and comparisons of marker distributions between different genotypes were assessed with the Chi-square method. All statistical parameters are indicated in Supplementary table 2.

## 3. Results

### 3.1. Ret and Brn3a co-expression changes dramatically during RGC development

We had previously demonstrated that ablation of Brn3a before the onset of its expression (E12), results in essentially complete loss of RGC types with small dense dendritic arbors (betta ON and OFF or “midget-like” RGCs), while other Brn3a^+^ RGC types are only modestly affected ^2,14,17^. ON and OFF β RGCs, alongside several other cell types, are labelled when sparse random recombination is induced in adult *Ret*^*CreERt2/WT*^; *Brn3a*^*CKOAP/WT*^ mice ^51^. To explore the time points at which Brn3a is required for betta RGC development, we induced random sparse recombination in *Ret*^*CreERt2/WT*^; *Brn3a*^*CKOAP/KO*^ and *Ret*^*CreERt2/WT*^; *Brn3a*^*CKOAP/WT*^ mice at E15, P0 and P22 to generate either isolated Ret^KO/WT^; Brn3a^KO/KO^ RGCs in a Ret^KO/WT^; Brn3a^KO/WT^ retinal background or Ret^KO/WT^; Brn3a^KO/WT^ RGCs in a Ret^KO/WT^; Brn3a^WT/WT^ retinal background (Figure 1 a, b, Figure 2 a – c1). In these experiments, the Brn3a gene dosages of labelled RGCs are different from the surrounding tissue. In order to study the effects of complete double heterozygosity of Ret and Brn3a (Ret^KO/WT^; Brn3a^KO/WT^) on RGC development, we induced random sparse recombination in either *Ret*^*CreERt2/WT*^; *Brn3a* ^*KO/WT*^; *Rosa26*^*AP/WT*^ or *Ret*^*CreERt2/WT*^; *Brn3a* ^*WT/WT*^; *Rosa26*^*AP/WT*^ mice. In these experiments, labelled RGCs and surrounding retina have the same genotype (either Ret^KO/WT^; Brn3a^KO/WT^ or Ret^KO/WT^; Brn3a^WT/WT^, Figure 1 c, d, Figure 6 a-a1).

As previously shown ^51^, Brn3a expression in adult retina predominantly intersects with Ret expression in five morphological RGC types – three monostratified (ON and OFFβ - “midget-like”, Figure 2 f-f2, and ON spiny, Figure 2 e-e2) and two bistratified (ON-OFF direction selective = ON-OFF-DS - Figure 4 a-a3, and Small Bistratified/Suppressed-by-contrast, henceforth SbC,, Figure 4 b-b3). In addition, isolated instances of several other monostratified cells, as well as a significant number of bistratified RGCs with recursive dendrites were observed (Figure 4, c-c3). A similar range of RGC types was observed when recombination was induced in *Ret*^*CreERt2/WT*^; *Brn3a*^*CKOAP/WT*^ mice at P0. However, the overall RGC type distribution changed dramatically when recombination was induced at E15. The ratio of monostratified:bistratified RGCs changed from ~33:66 % at P0 and P22 to 60:40 % at E15 (Figure 3 b-b2, Table 2). Two cell types, On-Alpha-Sustained = ONαS and Pixel Detectors (M5) (Figure 2 d-d2, g-g2), which are not observed in samples induced at P0 and P22, made up more than 50 % of monostratified cells (plots and pie-chart Figure 3 c-g, Table 2). Amongst E15-induced bistratified neurons, SbC morphologies were essentially missing, while three “novel” morphological types, not observed in the samples induced at P0 and P22 made up sizeable fractions of bistratified neurons (AT1, AT2, ON-Direction Selective = ON-DS, Figure 4 e-e3, g-g4, h-h4, Figure 5 a-f, Table 2). AT1 and AT2 are two unusual bistratified morphologies characterized by ON dendritic arbors which stratify in apposition to the GCL (normalized ID index is between ~0 and 0.2, where 0 is GCL level, Figure 5 a, b, left panels), while their OFF dendritic arbors laminate close to the INL (Figure 5 a, c, left panels). In the case of AT1, the OFF arbor is relatively simple and derives in most cases via a single branch straight from the cell body. However, AT2 bistratified neurons have a thicker OFF dendritic arbor (in z dimension) (Figure 5 a, c left plots), which can be occasionally resolved into two sub-arbors, creating the impression of a tri-stratified neuron (Figure 4 h-h4). Morphologies similar to AT2 were recovered by random sparse recombination in wild type retina (Badea et al 2004,^1^ Figure 15 a), and resemble cell type 85 in the serially reconstructed dataset in the Eyewire Museum (http//www.museum.eyewire.org). However, we are not aware of any instances of AT1 morphologies in either repositories or previous literature. Four of the five cell types which are unique to the Ret^CreERt2/WT^; Brn3a^AP/WT^ RGC population induced at E15 (ONαS, M5, AT1, AT2), were never previously observed in sparsely labelled Brn3a^AP/WT^ RGCs ^2,14,17,51^. In addition, the ON dendritic arbors of these 4 cell types are laminated within the IPL in close proximity to the GCL, a sublamina that is typically not labelled when dendritic arbors of the entire Brn3a^AP/WT^ RGC population are labelled by E9.5 recombination (^14,17,23^ and Figure 7a). The fifth, ON-DS, is, based on dendritic arbor areas and lamination, most likely the ON-DS RGC, which is known to occasionally have smaller branches co-stratifying with the OFF ChAT band, and has been recorded in early embryonic induced samples (^2,14,17^, Figure 4 e-e3, Figure 5 left side). Thus, it appears that a subset of E15 induced Ret^CreERt2/WT^; Brn3a^AP/WT^ RGCs constitute a distinct RGC subpopulation that typically does not usually express Brn3a or exhibits morphological differences from previously described Brn3a^+^ RGCs.

**Figure 4.**
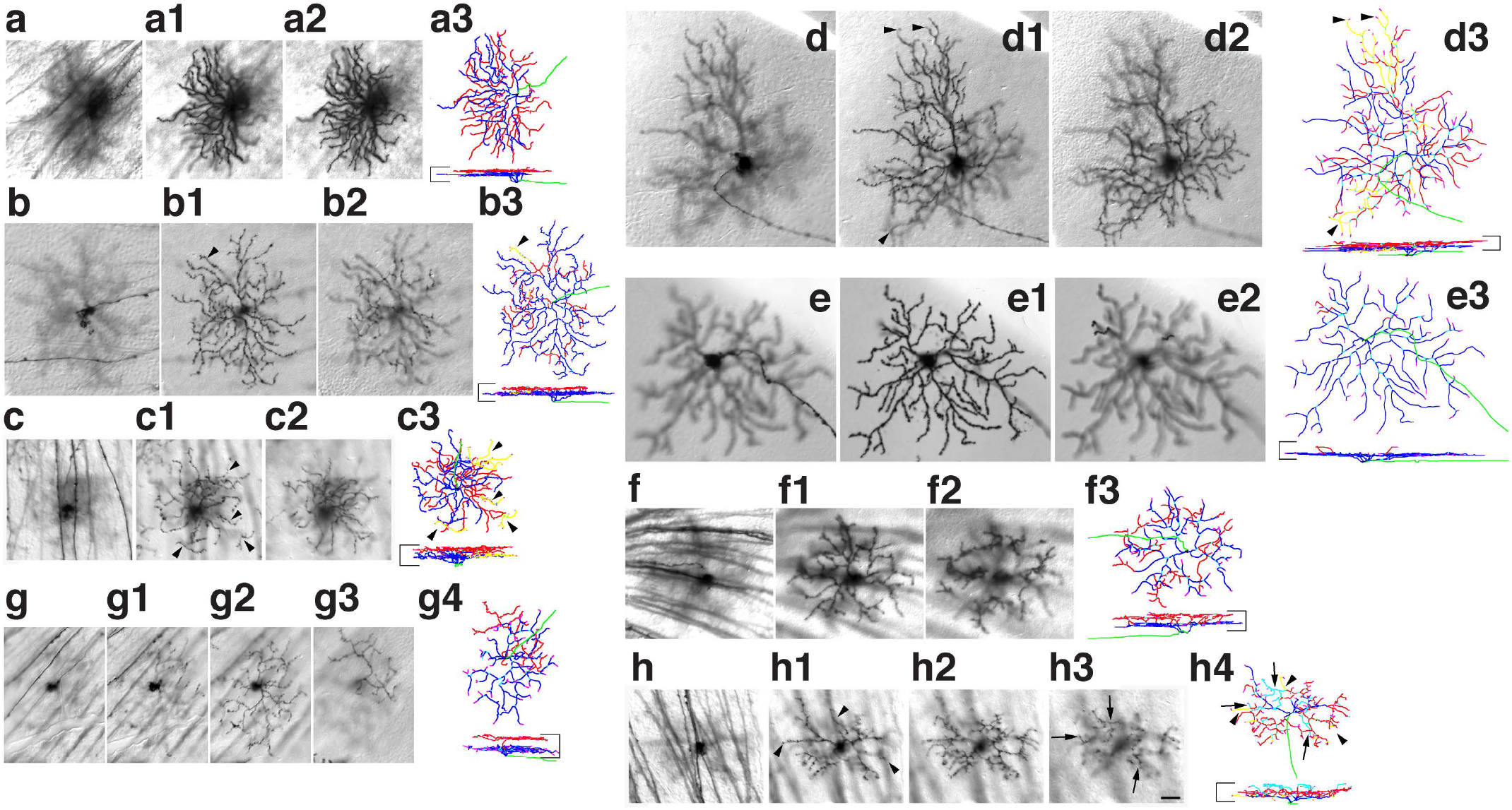
Bistratified RGCs in Ret^CreERt2^ Brn3a^CKOAP^ retinas. The most frequently encountered bistratified RGC morphologies were ON-OFF DS (a-a3), SbC (b-b3), Recursive (c-c3), Large bistratified (d-d3), ON-DS (e-e3), Brn3aKO-specific bistratified (f-f3), Abnormal type 1 (g-g4), and Abnormal type 2 (h-h4). For each cell focal planes through the cell body and axon (a, b, c, d, e, f, g, h), and the ON (a1, b1, c1, d1, e1, f1, g1-g2, h1) and OFF (a2, b2, c2, d2, e2, f2, g3, h2-h3) dendritic arbors are shown. Reconstructions, using the same conventions as in Figure 1, are shown in (a3, b3, c3, d3, e3, f3, g4, h4). Scale bar in (h3) and horizontal bars delineating the IPL in reconstructions are 25 μm. Axons are green, vitreal (ON) dendrites are blue, scleral (OFF) dendrites are red, second (scleral) sub-layer of OFF dendrites in Abnormal bistratified type 2 cell is cyan (arrows in h3-h4). Recursive dendrites laminating in the ON arbor but branching from the OFF arbor, are labeled in yellow in b3, c3, d3 and h4 (arrowheads in b1 and b3, c1 and c3, d1 and d3, and h1 and h4).

**Figure 5.**
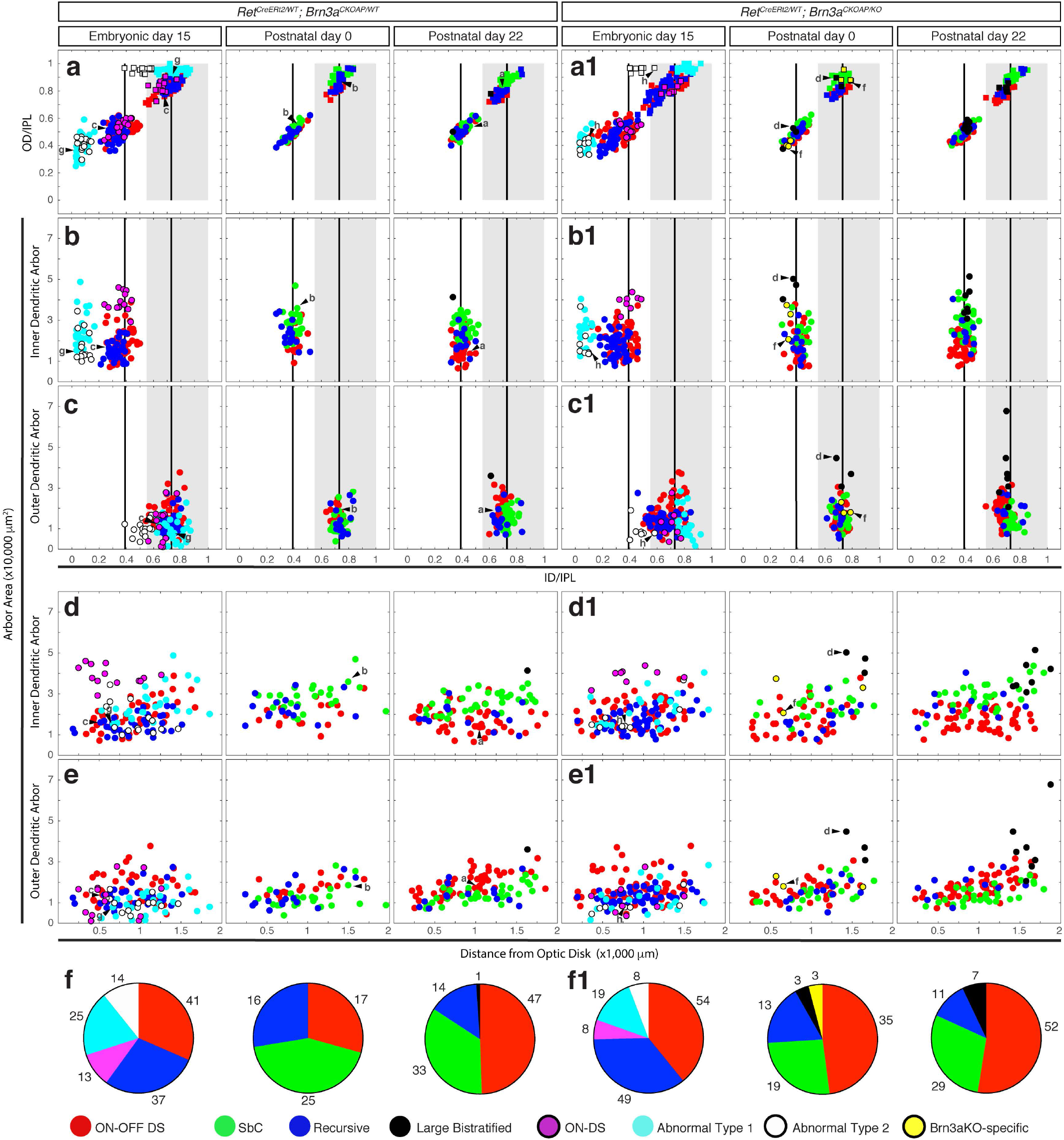
Morphological characterization of Ret^CreERt2/WT^; Brn3a^AP/WT^ and Ret^CreERt2/WT^; Brn3a^AP/KO^ bistratified RGCs. Scatter plots for morphological parameters of dendrites, defined as in Figure 2. (a-a1) Normalized ID (x) versus OD (y) measurement for either the inner (ON, circles) or outer (OFF, squares) dendritic arbors. (b-b1) Normalized ID of the inner arbor (x) versus inner arbor area (y). (c-c1) Normalized ID of the outer arbor (x) versus outer arbor area (y). (d-d1) Distance from optic disc to the cell body (x axis) versus inner arbor area (y). (e-e1) Distance from optic disc to the cell body (x axis) versus outer arbor area (y). The putative morphological types are color coded in the scatter plots and indicated at the bottom. Example cells from Figure 4 are indicated by arrowheads and letters corresponding to each cell in Figure 4. (f-f1) Pie-charts for proportions of different bistratified RGC subtypes in Ret^CreERt2/WT^; Brn3a^CKOAP/WT^ (f) and Ret^CreERt2/WT^; Brn3a^CKOAP/KO^ (f1) retinas. Morphological types are color coded as in (a-e1). Numbers in (f-f1) represent the total number of quantified RGCs for each genotype and subtype.

**Figure 6.**
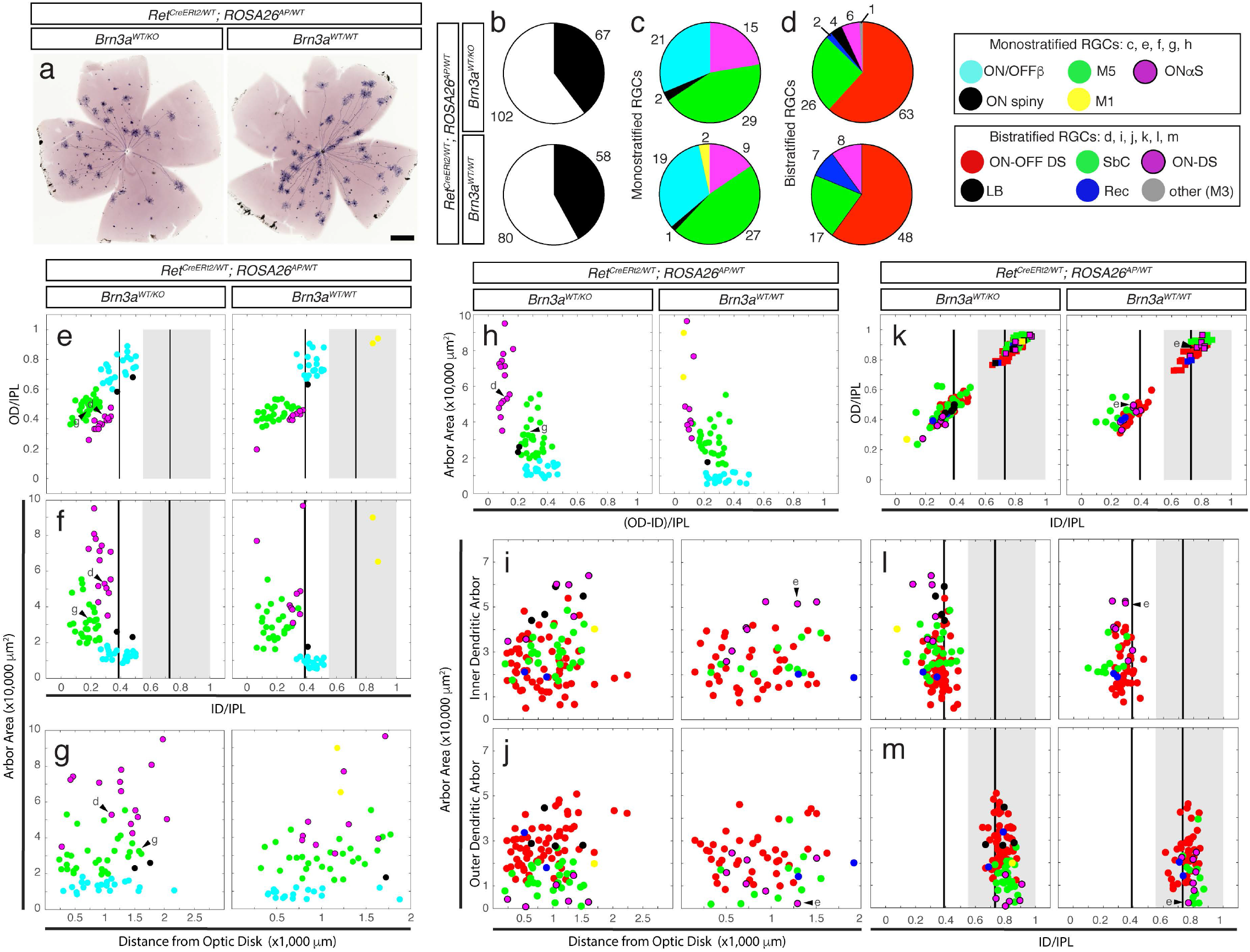
Morphological characterization of RGCs from *Ret*^*CreERt2/WT*^; *Brn3a*^*KO/WT*^; *ROSA26*^*AP/WT*^ and *Ret*^*CreERt2/WT*^; *Brn3a*^*WT/WT*^; *ROSA26*^*AP/WT*^ retinas. Cre was activated by i.p. administration of 250 μg 4-HT to mothers of the pups at gestational day 15 (E15 for pups). (a) Retinal flat mounts processed for AP histochemistry for *Brn3a*^*KO/WT*^ (left) and *Brn3a*^*WT/WT*^ (right) mice. Scale bar 500 μm. (b) Proportions of monostratified (black) to bistratified (white) RGCs in *Brn3a*^*KO/WT*^ (top) and *Brn3a* ^*WT/WT*^ (bottom) retinas. (c, d) Pie-charts for proportions of different monostratified (c) or bistratified (d) RGC types in *Brn3a*^*KO/WT*^ (top) and *Brn3a*^*WT/WT*^ (bottom) retinas. Numbers in (b-d) represent the total number of quantified RGCs for each genotype and category. Cell Numbers were derived from one or both retinas from at least three mice of each genotype. The color legends indicate monostratified types measured in scatter plots (e-h) and counted in pie charts (c), and bistratified types measured in (i-m) and counted in pie charts (d). (e-h) Scatter plots for parameters of monostratified RGCs, defined and presented as in Figure 2. (i-m) Scatter plots for parameters of bistratified RGCs. Example cells from Figures 2 and 4 are indicated by arrowheads and letters corresponding to each cell in Figure 2 – for monostratified RGCs, and in Figure 4 – for bistratified RGCs.

**Figure 7.**
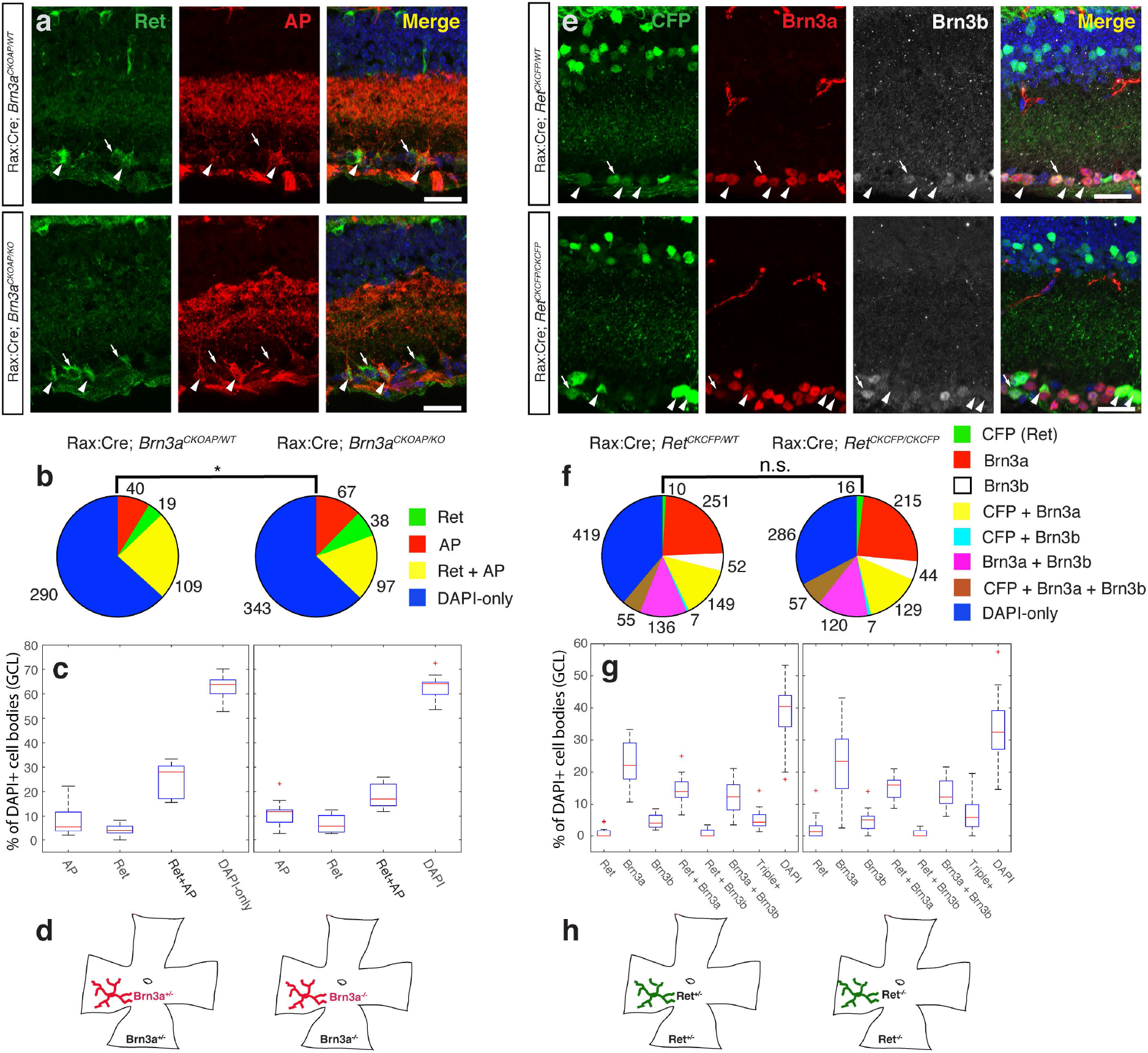
Effects of complete loss of function of Brn3a or Ret on RGCs. (a) Immunostaining of retina sections from Rax:Cre; *Brn3a*^*CKOAP/WT*^ (top row) and Rax:Cre; *Brn3a*^*CKOAP/KO*^ (bottom row) mice, using anti-Ret (green), anti-alkaline phosphatase (anti-AP, red) antibodies, and DAPI nuclear stain. Arrows indicate Ret-only-positive cells, arrowheads indicate Ret^+^AP^+^ double-labeled cells. (b) Pie-charts and (c) box-plots representing proportions of different cell categories according to the expression of Ret and AP in Rax:Cre; *Brn3a*^*CKOAP/WT*^ (left) and Rax:Cre; *Brn3a*^*CKOAP/KO*^ (right) animals. Total number of cells for each category are presented next to the pie-chart markers, and data spread for each category is shown in the boxplots, expressed as percent total DAPI positive cells. (d) RGCs (in red) and their respective retinas (in black) have matching genotypes in both Rax:Cre; Brn3a^CKOAP/WT^ (left) and Rax:Cre; Brn3a^CKOAP/KO^ (right) animals. (e) Immunostaining of sections through retinas from Rax:Cre; *Ret*^*CKCFP/WT*^ (top row, “Ret^KO/WT^”) and Rax:Cre; *Ret*^*CKCFP/CKCFP*^ (bottom row, “Ret^KO/KO^”) animals, using anti-GFP (staining CFP expressing cells, green) together with anti-Brn3a (red), anti-Brn3b (white) antibodies, and DAPI nuclear stain. Arrowheads indicate (CFP^+^Brn3a^+^) double-positive cells, and arrows indicate (CFP^+^Brn3a^+^Brn3b^+^) triple-labeled cells respectively. (f) Pie charts and (g) box-whiskers plots representing quantitation of immunostaining sections from Ret^KO/WT^ (left) and Ret^KO/KO^ (right) mice. Total number of cells for each category is presented next to the pie-chart markers. (h) CFP-positive RGCs (in green) and their respective retinas (in black) have matching genotypes in both Rax:Cre; *Ret*^*CKCFP/WT*^ (left) and Rax:Cre; Ret^CKCFP/CKCFP^ (right) animals. For each genotype in (a-c and e-g), sections from at least 3 different animals were stained and 9-20 images were quantified. For Box-Whisker plots, the red lines represent the median, the rectangles represent the interquartile interval, and the whiskers the full range of observations. Scale bar in (a) and (e) is 25 μm.

### 3.2. Loss of ON and OFF betta and ON spiny RGCs in Brn3a-cKO retinas after embryonic and perinatal Cre induction

We then analyzed subpopulations of Brn3a^KO/KO^ RGCs generated by inducing sparse recombination at E15, P0, and P22 (adult). When recombination was induced in the adult, type distribution of Ret^CreERt2/WT^; Brn3a^AP/KO^ (i.e. Ret^KO/WT^; Brn3a^KO/KO^) RGCs was indistinguishable from that of Ret^CreERt2/WT^; Brn3a^AP/WT^ (i.e. Ret^KO/WT^; Brn3a^KO/WT^ cells (Figure 3 b2, c-g1, Figure 5 a-f1, right-most pie charts and scatter plots, Table 2). Monostratified:Bistratified ratios were nearly identical (Figure 3 b2, Table 2), and the frequencies of individual mono- and bistratified RGC types was not significantly different (Figure 3 g-g1, Figure 5 f-f1, right-most pie charts and scatter plots, Table 3). However, when recombination was induced at P0 - resulting in random sparse Brn3a loss of function soon after birth - the distributions of Ret^CreERt2/WT^; Brn3a^AP/KO^ and Ret^CreERt2/WT^; Brn3a^AP/WT^ RGC types showed significant differences (Figure 3 b1, c-g1, Table 2, Table 3). The relative abundance of ON and OFFβ RGCs decreased considerably, and ON-Spiny neurons were completely missing (Figure 3 c-g1, middle plots and pie-charts), resulting in an overall decrease in monostratified RGC morphologies (Figure 3 b1, Table 2). Additionally, we identified amongst Ret^CreERt2/WT^; Brn3a^AP/KO^ RGCs induced at either P22 or P0 a minor subpopulation of RGCs with morphologies reminiscent of betta RGCs, but exhibiting somewhat larger areas and sparser dendritic arbors (Brn3aKO-specific betta RGC = Brn3aKO-β, Figure 2 h-h2, Figure 3c1-g1, middle and right scatter plots and pie charts).

RGCs with ON and OFF β and ON Spiny morphologies are also underrepresented amongst Ret^CreERt2/WT^; Brn3a^AP/KO^ RGCs induced at E15, when compared to Ret^CreERt2/WT^; Brn3a^AP/WT^ control RGCs (Figure 3 c-g1, left plots and pie-charts, Table 2) while the relative ratios of ONαS and Pixel Detector (M5) RGCs are not significantly affected by complete loss of Brn3a (Figure 3 c-g1, Table 2). These shifts in cell type distribution result in a net reduction of the mono:bistratified ratio in Ret^CreERt2/WT^; Brn3a^AP/KO^ RGCs induced at E15 compared to controls. (Figure 3 b, g-g1 left pie-charts, Table 2).

### 3.3. Effect of Brn3a loss of function on bistratified RGCs

In our previous work, we had reported a moderate effect of Brn3a ablation on bistratified RGC morphology. Here we use sparse random recombination in the *Ret*^*CreERt2*^; *Brn3a*^*CKOAP*^ intersection to study the effect of Brn3a ablation on bistratified RGCs at several developmental stages. Bistratified Ret^CreERt2/WT^; Brn3a^AP/WT^ and Ret^CreERt2/WT^; Brn3a^AP/KO^ RGC types did not differ significantly when sparse random recombination was induced at either P0 or P22 (Figure 5, a-f1, middle and right plots and pie-charts, Table 2). The majority of bistratified RGCs labeled belonged to the ON-OFF DS, SbC, and Recursive Bistratified types. A few examples of Large Bistratified RGCs (LB, Figure 4 d-d3) were also observed specifically amongst P0 and P22 Ret^CreERt2/WT^; Brn3a^AP/KO^ RGCs (Figure 5 a-f1, right plots and pie-charts). In addition, in P0 induced recombinations, a few instances of a Brn3aKO-specific subpopulation were recovered. These cells exhibit a simplified ON arbor from which relatively simple straight branches descend and form small tufts into the OFF lamina (Figure 4 f-f3, Figure 5 a1-e1 middle plots), and resemble the ones previously described in early sparse recombination experiments ^2^. The AT1 and AT2 morphologies are observed in the dataset induced at E15 in both Ret^CreERt2/WT^; Brn3a^AP/WT^ and Ret^CreERt2/WT^; Brn3a^AP/KO^ RGCs with comparable frequencies (Figure 5, f-f1, left pie-charts, Table 2).

### 3.4. Early Complete (Ret^KO/WT^; Brn3a^KO/WT^) double heterozygosity does not affect RGC type distribution

Ret expression in the retina changes significantly between embryonic, postnatal and adult stages of development ^51^. We had previously described the RGC type distribution in *Ret*^*CreERt2/WT*^; *ROSA26*^*AP/WT*^ mice in which random sparse recombination was induced at P14 and adult ^51^. Is the dramatic shift in Ret^+^ Brn3a^+^ RGC types observed in E15 random sparse recombination due to developmental differences of expression intrinsic to the Ret locus? Alternatively, are the shifts in RGC types due to a genetic interaction between Brn3a and Ret in the double heterozygote mice? To answer these questions, we induced random sparse recombination in *Ret*^*CreERt2/WT*^; *Brn3a*^*KO/WT*^; *ROSA26*^*AP/WT*^ and *Ret*^*CreERt2/WT*^; *Brn3a*^*WT/WT*^; *ROSA26*^*AP/WT*^ mice at E15 and analyzed RGC type distribution in adult (>P60) mice (Figure 1 c, d, Figure 6 a). The ratio between mono- and bistratified RGCs is similar in complete *Brn3a*^*WT/WT*^ and complete *Brn3a*^*KO/WT*^ retinas (Figure 6 b) and the distribution of monostratified RGC types induced at E15 in *Ret*^*CreERt2/WT*^; *ROSA26*^*AP/WT*^ mice was not affected by Brn3a dosage and resembled the RGC types identified in *Ret*^*CreERt2/WT*^; *Brn3a*^*CKOAP/WT*^ mice induced at the same age, with small variations in M5 and betta cell numbers (Figure 6 c, e-h, Table 2). However, bistratified RGCs induced at E15 in either *Ret*^*CreERt2/WT*^; *Brn3a*^*WT/WT*^; *ROSA26*^*AP/WT*^ or *Ret*^*CreERt2/WT*^; *Brn3a*^*KO/WT*^; *ROSA26*^*AP/WT*^ backgrounds were restricted to the previously described major types (ON-OFF DS, SbC, Recursive, ON-DS and LB), and no instances of AT1 and AT2 RGCs were observed (Figure 6 d, i-m). This distribution is consistent with the one seen for *Ret*^*CreERt2/WT*^; *Brn3a*^*CKOAP/WT*^ or *Ret*^*CreERt2/WT*^; *ROSA26*^*AP/WT*^ mice upon adult inductions. Thus, complete heterozygous Brn3a loss did not affect the pattern of Ret expression at E15. Moreover, the expression profile of Ret amongst RGC types, as measured by induction of *Ret*^*CreERt2/WT*^ in the background of the *ROSA26*^*AP/WT*^ allele, does not change dramatically from E15 to adult (compare Figure 6 h, k to Parmhans 2018, ^51^, Figures 6 h and 7 c-d).

### 3.5. Regulatory crosstalk between Ret and Brn3a

The distinct effects of global versus sparse random manipulation of Brn3a dosage on RGC type distribution prompted us to ask whether Ret and Brn3a regulate each other at transcriptional level. We therefore checked for co-expression of Ret protein and the Alkaline phosphatase (AP) reporter in full-retina Brn3a-heterozygote (*Rax:Cre; Brn3a*^*CKOAP/WT*^) and full-retina Brn3a-knockout (*Rax:Cre; Brn3a*^*CKOAP/KO*^, Figure 7 a-d) sections. The distribution of Ret^+^, AP^+^ and Ret^+^AP^+^ (double-positive) cells is similar regardless of Brn3a dosage, with a modest (statistically insignificant) decrease of double-positive cells in *Rax:Cre; Brn3a*^*CKOAP/KO*^ (Figure 7 b, c, Supplementary Table 2). Nevertheless, the overall shift from AP^+^Ret^+^ double positive cells to single AP^+^ or Ret^+^ positive cells in Brn3a^KO^ retinas is statistically significant (χ^2^ ChiStat = 10.81, p = 0.013) potentially suggesting that Brn3a controls Ret in a subset of RGC types.

We then asked whether Ret signaling can regulate Brn3a or Brn3b transcription. Co-expression of the conditional knock-in reporter CFP with either Brn3a or Brn3b was compared in full-retina Ret-heterozygotes (*Rax:Cre; Ret*^*CKCFP/WT*^) and full-retina Ret-knockouts (*Rax:Cre Ret*^*CKCFP/CKCFP*^, Figure 7 e-h). In this line, the CFP reporter is expressed from the Ret locus after the removal of the Ret cDNA by Cre recombination ^47^. We observed RGCs expressing Ret either alone or in combination with Brn3a, Brn3b or both. The distribution of single, double and triple labelled cells is conserved in the two backgrounds (Figure 7 f, g, Supplementary Table 2), suggesting that Ret is not required for Brn3a or Brn3b expression in RGCs. Thus it is unlikely that the genetic interaction observed in sparsely recombined RGCs is mediated by reciprocal transcriptional control of Ret and Brn3a.

### 3.6. Does Brn3a modulate GDNF ligand signaling to RGCs by regulating GFRα Ret co-receptors?

We next asked whether GFRα Ret co-receptors are expressed in Brn3a^+^ RGCs and/or regulated by Brn3a. Data from a deep sequencing analysis screen of Brn3a transcriptional targets expressed in RGCs ^9,23^ shows that Ret and GFRα’s are expressed in RGCs at E15 and P3 (Supplementary Figure 1). While Ret expression levels are relatively high (some 20 FPKM), the co-receptors are expressed at much lower levels. The major GFRα1 transcript is mostly expressed in Brn3b^AP^ RGCs at both E15 and P3 (around 5 FPKM), and somewhat less in Brn3a^AP/WT^ RGCs at P3 (1.2 FPKM), and its expression is nearly doubled in Brn3a^AP/KO^ RGCs, suggesting negative regulation by Brn3a. GFRα2 is homogeneously expressed across all Brn3^AP^ RGCs at both E15 and P3 (about 4-6 FPKM), but is not regulated by Brn3a. GFRα3 is and GFRα4 are expressed at less than 1 FPKM in P3 Brn3^AP^ RGCs and do not appear regulated by either Brn3 transcription factor (Sajgo 2017 and Supplementary Figure 1). Since adult GFRα expression had been previously reported in RGCs ^48^, we stained adult *Rax:Cre; Brn3a*^*CKOAP/WT*^ and *Rax:Cre; Brn3a*^*CKOAP/KO*^ retina sections with anti-GFRα1, GFRα2 and GFRα3 antibodies (Figure 8, Supplementary Table 2). When comparing *Brn3a*^*AP/WT*^ to *Brn3a*^*AP/KO*^ retinas (Figure 8, c-d, g-h, k-l Supplementary Table 2), loss of Brn3a results in significant increases in GFRα1^+^ - GFRα3^+^ GCL cells (18.8 to 25, 25.8 to 33.33 and 38.2 to 54 % DAPI^+^ cells in GCL, respectively). Consistent with previous reports, Brn3a^AP^ RGCs numbers are reduced as a result of Brn3a ablation (26 to 18.8, 20.18 to 9 and 4.35 to 2.4 % DAPI^+^ cells in GCL, respectively), however GFRα1^+^ Brn3a^AP^ and GFRα2^+^ Brn3a^AP^ double positive ratios are not significantly affected (GFRα1^+^ Brn3a^AP^: 7.5 vs. 11.7 and GFRα2^+^ Brn3a^AP^: 12.7 vs 11.3 % DAPI^+^ cells in GCL). There is also a sizable but statistically insignificant decrease in GFRα3^+^ Brn3a^AP^ RGCs. Overall, the partial overlap between all three GFRα receptors and Brn3a^AP^ is significantly altered by Brn3a ablation (Figure 8 c, g, k), resulting in a shift away from Brn3a^AP^ and towards GFRα expression. This shift could be caused by a fate change of Brn3a^AP^ RGCs towards GFRα^+^ RGCs in *Brn3a*^*AP/KO*^ retinas, since the ratios of GFRα^+^ Brn3a^AP^ RGCs are not significantly changed. Interestingly, the antibody staining for both GFRα1^+^ and GFRα2^+^ reveals increased dendritic arbor labelling in close proximity to the GCL, suggesting that most GFRα1^+^ and GFRα2^+^ RGCs are laminating in the sublamina which is populated by the unusual Ret^CreERt2/WT^; Brn3a^AP/WT^ or Ret^CreERt2/WT^; Brn3a^AP/KO^ RGC types induced at E15 (ONαS, M5, AT1, AT2).

**Figure 8.**
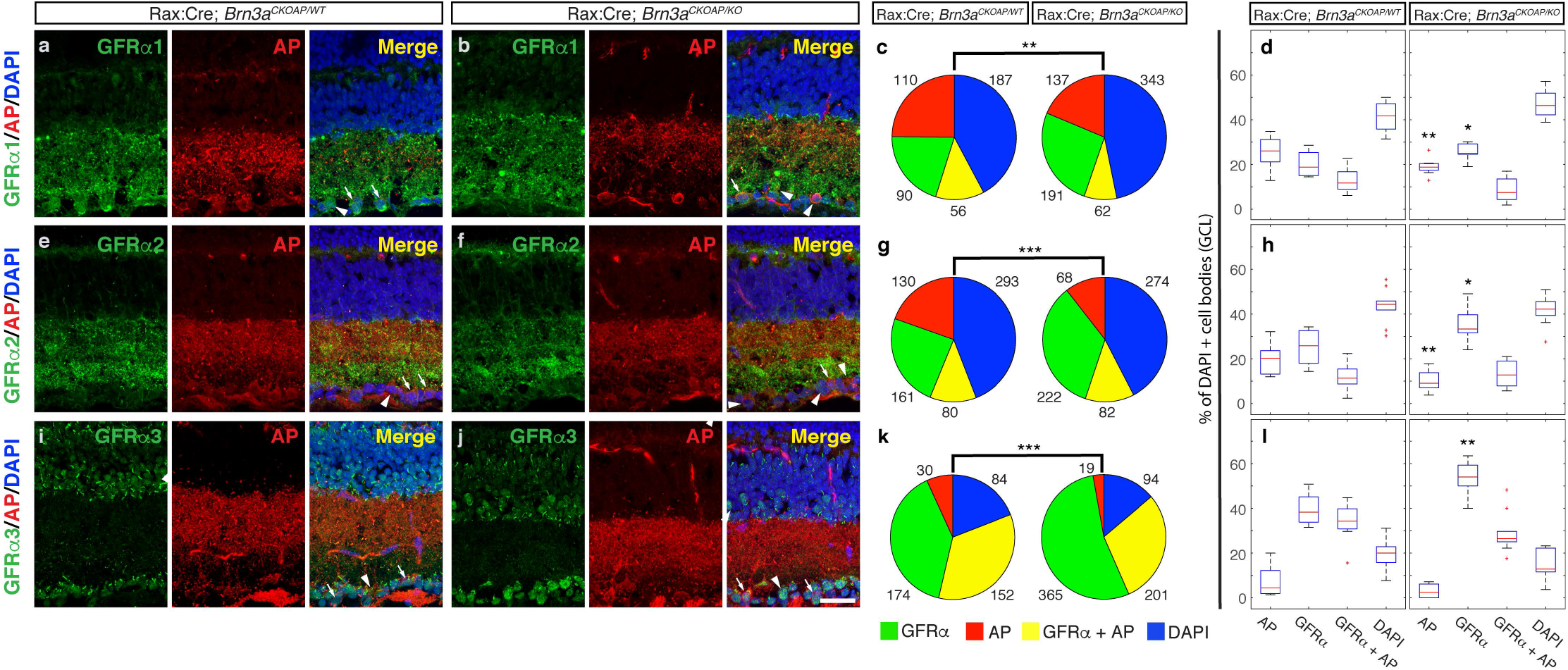
Full retinal Brn3a KO significantly affects distribution of GFRα Ret co-receptors in RGCs. Retinal sections from Rax:Cre; *Brn3a*^*CKOAP/WT*^ (a,e,i) and Rax:Cre; *Brn3a*^*CKOAP/KO*^ (b, f, j) mice, were stained using antibodies against GFRα1 (a-b), GFRα2 (e-f) or GFRα3 (i-j) (green, left panels) in conjunction with AP (red, middle panels) and DAPI (right panels show three-color merged image). Arrowheads indicate single labelled cells (AP or GFRα) and arrows point to double labeled cells (GFRα ^+^AP^+^). Total numbers of quantified cells are represented as pie charts in c, g and k and indicated next to each sector. Significance levels for the comparison between the distributions for Rax:Cre; *Brn3a*^*CKOAP/WT*^ (left) and Rax:Cre; *Brn3a*^*CKOAP/ KO*^ mice are calculated using the Chi-square statistic. Chi statistic, p-values and degrees of freedom are indicated in Supplementary table 2. Box-plots representing proportions of different cell categories according to the expression of each GFRα and AP are shown in d, h an l, expressed as percent total DAPI positive cells in the GCL. Significance levels for differences between Rax:Cre; *Brn3a*^*CKOAP/WT*^ (left) and Rax:Cre; *Brn3a*^*CKOAP/KO*^ (right) retinas were calculated using Kolmogorov-Smirnoff 2 test, and number of quantitated cells, sections and mice for each genotype are indicated in Supplementary table 2. For Box-Whisker plots, the red lines represent the median, the rectangles represent the interquartile interval, and the whiskers the full range of observations. Outliers are indicated by red crosses. Scale bar in (j) is 25 μm.

**Figure 9.**
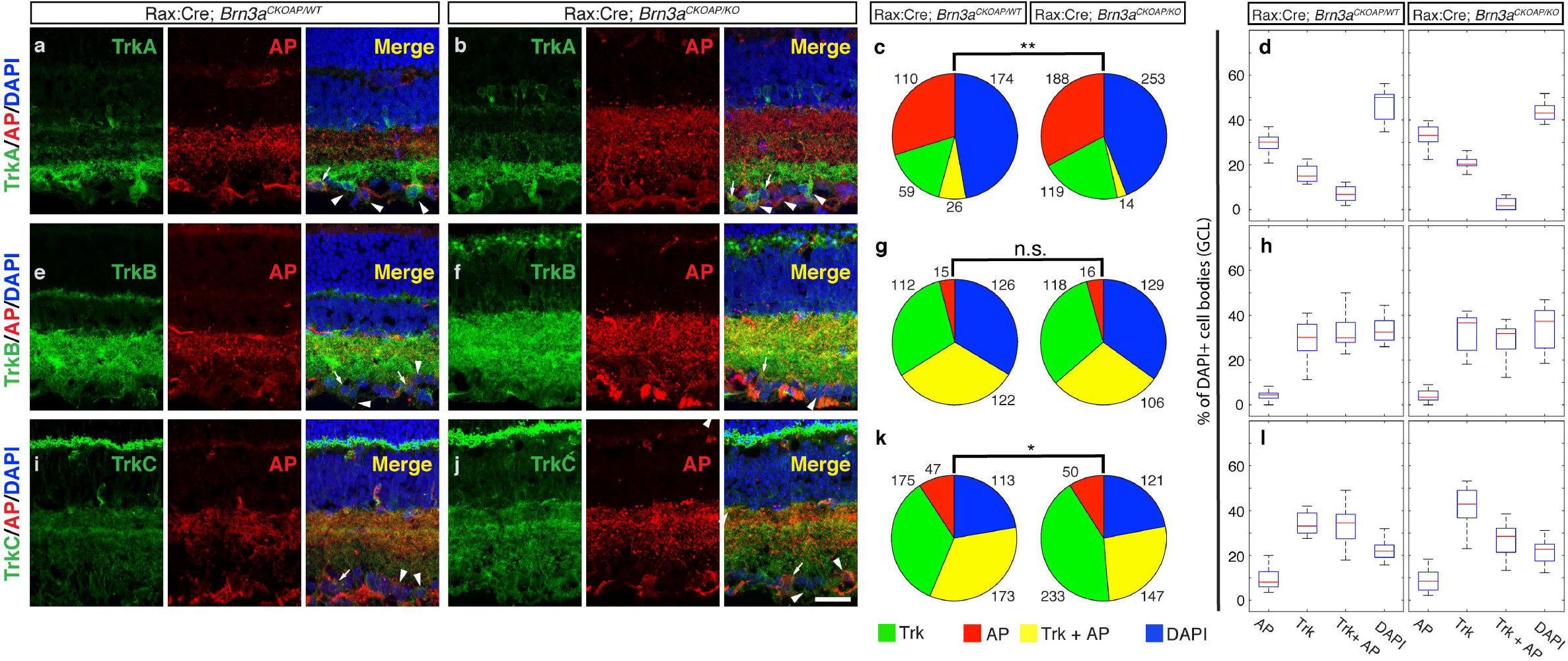
Full retinal Brn3a KO mildly affects distribution of Trk receptors in RGCs. Retinal sections from Rax:Cre; *Brn3a*^*CKOAP/WT*^ (a,e,i) and Rax:Cre; *Brn3a*^*CKOAP/KO*^ (b, f, j) mice, were stained using antibodies against TrkA (a-b), TrkB (e-f) or TrkC (i-j) (green, left panels) in conjunction with AP (red, middle panels) and DAPI (right panels show three-color merged image). Arrowheads indicate single labelled cells (AP or Trk) and arrows point to double labeled cells (Trk^+^AP^+^). Total numbers of quantified cells are represented as pie charts in c, g and k and indicated next to each sector. Significance levels for the comparison between the distributions for Rax:Cre; *Brn3a*^*CKOAP/WT*^ (left) and Rax:Cre; *Brn3a*^*CKOAP/KO*^ mice are calculated using the Chi-square statistic. Chi statistic, p-values and degrees of freedom are indicated in Supplementary Table 2. Box-plots representing proportions of different cell categories according to the expression of each Trk receptor (A, B or C) and AP are shown in d, h an l, expressed as percent total DAPI positive cells in the GCL. Significance levels for differences between Rax:Cre; *Brn3a*^*CKOAP/WT*^ (left) and Rax:Cre; *Brn3a*^*CKOAP/KO*^ (right) retinas were calculated using Kolmogorov-Smirnoff 2 test, and number of quantitated cells, sections and mice for each genotype are indicated in Supplementary table 2. There were no significant shifts in any of the single or double labelled populations. For Box-Whisker plots, the red lines represent the median, the rectangles represent the interquartile interval, and the whiskers the full range of observations. Scale bar in (j) is 25 μm.

### 3.7. Does Brn3a transcriptionally regulate Trk family neurotrophic receptors in RGCs?

Brn3a regulation of Trk neurotrophin receptors is believed to play a major role in cell type specification of projection sensory neurons of the somatosensory (DRG and TGG), auditory and vestibular pathways, and the GDNF - GFRα and NGF – Trk neurotrophic signaling axes interact in cell type specification^25,26,56^. We therefore asked whether Trk receptor expression in RGCs is regulated by Brn3a. Our RNAseq data predicted that all three Trk receptors (TrkA/Ntrk1, TrkB/Ntrk2 and TrkC/Ntrk3) and p75/NGFr are expressed in RGCs at E15 and P3, and that Brn3a is positively regulating TrkA and TrkC and negatively regulating TrkB (^9,23^ and Supplementary Figure 2), while TrkB expression in adult mouse RGCs had been previously reported^57^. Using antibody staining in the adult retina, we find that TrkB is expressed in a large fraction of GCL cells, and a majority of Brn3a^AP^ RGCs in both *Brn3a*^*AP/WT*^ or *Brn3a*^*AP/KO*^ retinas (Figure 9 e-h, Supplementary Table 2). In contrast, TrkA is partially co-expressed with Brn3a in a small fraction of RGCs, and the fraction of TrkA^+^ Brn3a^AP^ RGCs is somewhat reduced in *Brn3a*^*AP/KO*^ retinas (from 6.8 to 1.8 % DAPI^+^ cells in GCL, Figure 9 a – d, Supplementary Table 2). A majority of Brn3a^AP^ RGCs expressed TrkC, and the number of TrkC^+^ Brn3a^AP^ RGCs was mildly reduced by Brn3a ablation (from 34.6 to 28.5 % DAPI^+^ cells in GCL, Figure 9 i-l, Supplementary Table 2). While none of the individual cell population changes were statistically significant using the KS2 test, TrkA and TrkC vs. Brn3a^AP^ populations were significantly shifted in *Brn3a*^*AP/WT*^ vs *Brn3a*^*AP/KO*^ retinas, as judged by the Chi-Square distribution test. These losses in TrkA^+^ Brn3a^AP^ and TrkC^+^ Brn3a^AP^ RGCs may be due either to direct transcriptional regulation of the two Trk receptors by Brn3a or by the loss of specific Brn3a^AP^ RGC subpopulations due to Brn3a ablation. It is worth pointing out that TrkA dendritic arbors were distributed in three sharp lamina across the IPL, with the most intense one being apposed against the GCL, as seen for GFRα1^+^ and GFRα2^+^, while TrkC exhibited lamination in the OFF sublaminae of the IPL. All three Trk receptors are expressed in the GCL and the proximal INL, suggesting expression in Amacrine cells in addition to RGCs.

## 4. Discussion

Our results show that altering the dosage of Brn3a in a sparse mosaic fashion early (E15) in the development of Ret heterozygote (Ret^CreERt2/WT^) RGCs can change the cell type distribution and/or morphologies of heterozygote (Brn3a^AP/WT^) *<u>and</u>* knockout (Brn3a^AP/KO^) RGCs. RGCs are not affected when Brn3a is removed in the adult, and mildly affected by Brn3a removal immediately after birth. E15 or P0 removal of both copies of Brn3a results in dramatic losses in midget and ON spiny RGCs, suggesting a significant role for Brn3a in the development of these cell populations. Germline double heterozygosity (Ret^CreERt2/WT^; Brn3a^KO/WT^) does not phenocopy the results obtained with the mosaic heterozygotes and results in normal RGC type specification. Immunohistochemical evidence collected in whole retina knockouts of either Ret or Brn3a shows a modest transcriptional control of Ret by Brn3a while neither Brn3a nor Brn3b are affected by complete retinal loss of Ret. However full retinal loss of Brn3a results in significant shifts in expression of Ret co-receptors GFRα1-3 in the GCL and mildly reduces neurotrophin receptors TrkA and TrkC expression in Brn3a^AP^ RGCs. Thus, the simplest explanation of our data is that Ret and Brn3a participate in parallel developmental pathways that converge onto RGC type specification at the early stages of postmitotic development, and potentially use a competitive mechanism based on gene dosage. The range of RGC types reported in this study was largely similar with previous reports, and equivalencies to previously reported anatomies, including a serial EM dataset (EyeWire museum)^4^ are provided in table 1.

**Table 1.** RGC subtypes and their possible matches from EyeWire Museum For each type we provide the closest correspondence in the EyeWire museum and physiology or anatomy literature, if available.

| RGC subtype | #-s of RGCs from EyeWire Museum | References |
| --- | --- | --- |
| *Monostratified RGCs* |  |  |
| ON alpha sustained | 8w (ONαS) | 90-92 |
| M5 | 8n, 9n | 93-96 |
| ON spiny | 6sn, | 97-100 |
| ON betta | 63 (F-mini<sup>On</sup>), 5ti, 5si | 98,100,101 |
| OFF betta | 2an (F-mini<sup>Off</sup>) | 98,100,101 |
| OFF transient | 4i, 4on, 4ow, 3o | 90,91 |
| OFF sustained | 1no, 1ni, 1wt | 90,91 |
| *Bistratified RGCs* |  |  |
| ON-OFF DS | 37c, 37d, 37r, 37v (On-Off DS) | 102-105 |
| Suppressed-by-contrast | 7o(tOn DS), 81i, 82n | 106-108 |
| Recursive | 72, 73 | 2 |
| Large Bistratified | n/a | 2,51 |
| ON - DS | 7id, 7ir, 7iv (sOn DS) | 109,110 |
| Abnormal type 1 | n/a | n/a |
| Tristratified (Abnormal Type 2) | 85 | 1 |

**Table 2.**
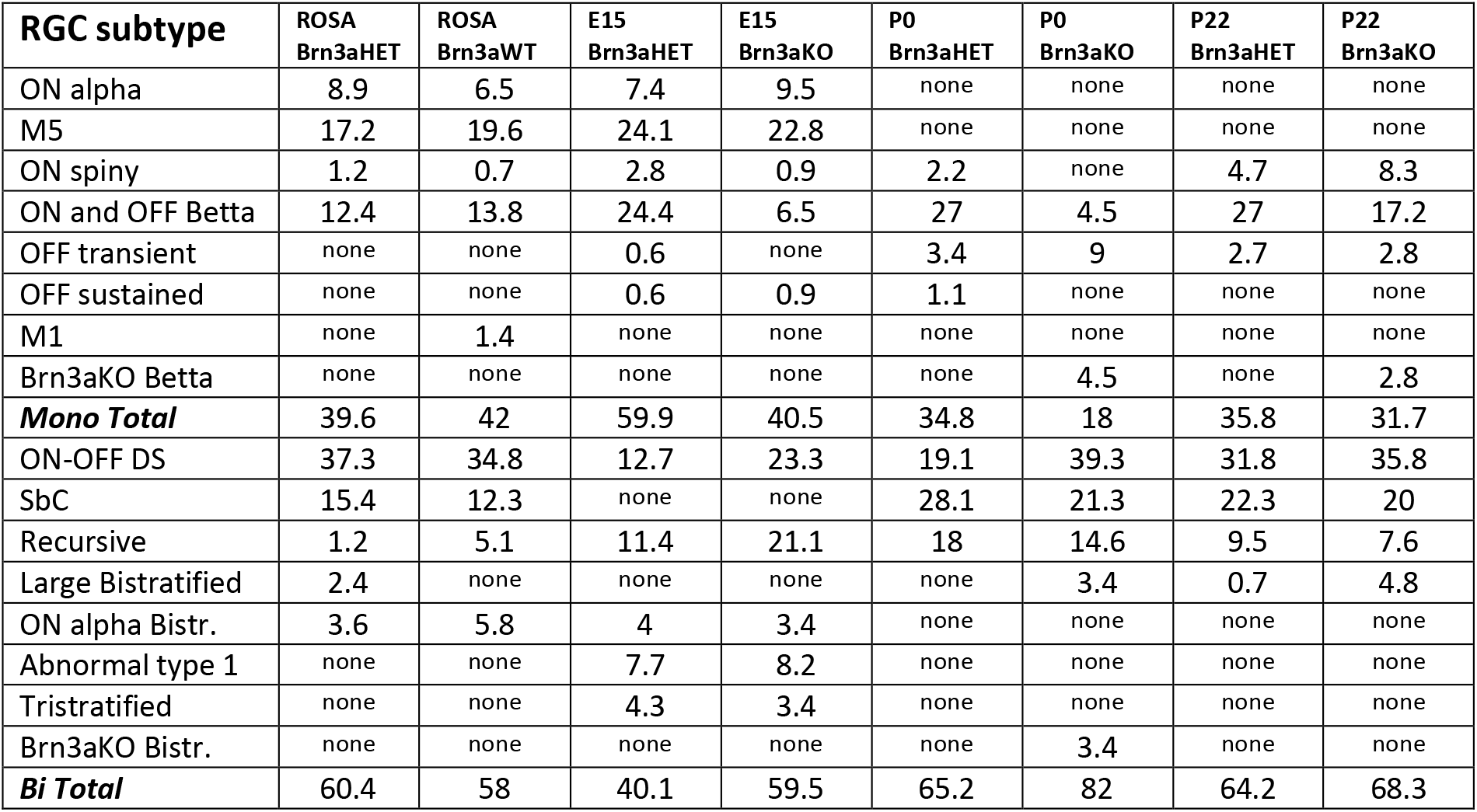
RGC subtype frequencies in different experimental ages and groups (relative to the total number of RGCs in an experimental group)

### Brn3a requirement in early versus late RGC development

Brn3a is required for the development of ON and OFF β RGCs and some bistratified RGC types, and Brn3a KO animals have a net RGC loss of about 30 % ^2,14,17,51^. Ablating either Brn3a, Brn3b or both in adult mice does not affect RGC numbers up to six months post ablation ^58^. We now show that adult ablation of Brn3a does not significantly alter the cell type distribution of Ret^+^ Brn3a^+^ RGCs, while complete loss of Brn3a at P0 and E15 produces a significant overall shift in RGC type distribution between Ret^CreERt2/WT^ Brn3a^AP/WT^ and Ret^CreERt2/WT^ Brn3a^AP/KO^ RGCs, largely based on the dramatic loss of ON and OFF β RGCs (ratios in Het vs. KO are 27% vs. 17.2% at P22, 27% vs. 4.5 % at P0 and 24.4% vs. 6.5% at E15). Thus, Brn3a is required for the development of these cells but not for maintenance in the adult. In P0 and P22 induced Brn3a^AP/KO^ RGCs we also detected an unusual dendritic arbor morphology reminiscent of ON betta RGCs, but distinguishable by larger areas and less dense dendritic coverage potentially representing abnormal ON betta RGCs resulting from relatively late loss of Brn3a. ON spiny RGCs are also selectively reduced in E15 and P0 but not adult inductions, suggesting an early dependency on Brn3a. Furthermore, two types of bistratified RGCs (LB and Brn3aKO – bistratifieds) are observed specifically in P0 and P22 induced Brn3a^AP/KO^ RGCs. LB (Large Bistratifed) RGCs have been previously seen in WT circumstances ^2,17,51^, especially in Brn3b^+^ RGCs, and therefore could represent another shift of cell type specificity, induced only in Ret^CreERt2/WT^ Brn3a^AP/KO^ RGCs. However, the simplified arbors of Brn3aKO-bistratifieds point to a developmental defect as a result of Brn3a loss, and is independent of Ret, since they were observed in Brn3a ablations using other Cre drivers ^2,14^.

### Ret and Brn3a: Genetic interaction versus developmental dynamic shift of expression

Is it possible that the shift in RGC morphologies seen upon embryonic recombination reflects the dynamic expression pattern of Ret or Brn3a in RGCs? The dynamic expression pattern of Ret was shown to be important for specification of subpopulations of DRG neurons ^44,45^ and we have documented the dynamic expression of Ret in RGCs, Horizontal cells and Amacrines during embryonic and postnatal retinal development. However, the distribution of RGC types in *Ret*^*CreERt2/WT*^; *ROSA26*^*AP/WT*^ mice seems to be relatively stable from E15 through P14 to adult, (data in figure 7 in this study and figures 6 – 7 in ^51^), arguing that the expression profile of Ret^CreERt2^ is relatively unchanged during RGC development. The cell type distribution is collected and analyzed in adult mice in all experiments, using AP expressed under the control of the Brn3a locus (Brn3a^AP^), thus reflecting only the adult expression profile of Brn3a.

The RGC morphologies observed when recombination is induced in adult *Ret*^*CreERt2/WT*^; *Brn3a*^*CKOAP/WT*^ mice are in agreement with our previously published data regarding Brn3a expression in various RGC types. In previous studies, recombination was achieved using either transgenic elements (Pax6α:Cre or Rax:Cre) that are activated at E9.5 to 10.5, or sparse random recombination induced using alleles with no known biological effects or preferential expression patterns, under the control of the ROSA26 locus or CAG promoter, induced at a variety of ages, from E8 to adult. The repertoire of RGC types positive for Brn3a revealed by sparse labelling using general promoters (ROSA26, Pax6α:Cre) shows a dendrite lamination pattern typically excluding the innermost 20-30 % of the IPL. This pattern is confirmed by immunofluorescent staining in sections where the totality of Brn3a^AP^ RGC dendrites is revealed using whole retina Cre drivers such as Pax6α:Cre and Rax:Cre (^14,17,23^ and Figure 7a in this study). It is therefore likely that P0 and adult-induced Ret^CreERt2/WT^; Brn3a^AP/WT^ RGCs accurately reflect the expression overlap of Ret and Brn3a in RGCs.

Overall, five cell types are unique to the E15 induced Ret^CreERt2/WT^; Brn3a^AP/WT^ RGCs population as contrasted to P0/adult induced RGCs: ONαS, M5, ON-DS, AT1 and AT2. They make up a sizeable fraction (47%) of all E15 induced RGCs. Of these, only ON-DS cells are known to be Brn3a positive in the adult and are also present in the general E15 Ret RGC expression profile (*Ret*^*CreRt2*^; *Rosa26*^*iAP*^ data), and thus could be explained by a shift of expression of Ret between E15 and P0/adult. The other four cell types have not been previously reported to be Brn3a positive. Furthermore, while ONαS, M5, and ON-DS cells are present in E15 and adult induced *Ret*^*CreRt2*^; *Rosa26*^*iAP*^ mice, AT1 and AT2 are completely missing from these data sets. Since ONαS and M5 can be reliably matched to Ret^+^ RGC types, their appearance in E15 induced Ret-Brn3a double heterozygotes could be interpreted as an induction of Brn3a expression in these cell types, or a cell type shift caused by the genetic interaction between Ret and Brn3a. In contrast AT2 and AT1 could be derived from a faulty cell type specification decision, separating them from the normal Ret^+^Brn3a^+^ fate and resulted from the combined heterozygote loss of Brn3a and Ret. AT2 is tentatively matched morphologically to cell type 85 in the EyeWire museum, and we described one instance in a general RGC description in 2004^1^, and AT1 does appear to be a novel morphology. It is therefore possible that AT1 and AT2 morphologies result from developmental changes in SbC cells, which are absent from E15 specified Ret^CreERt2^ Brn3a^AP^ RGCs, but are present in large numbers in E15 induced Ret^CreRt2^; Rosa26^iAP^ RGCs. We conclude that labelling of ONα, M5, AT1 and AT2 RGCs in Ret^CreERt2/WT^; Brn3a^AP/WT^ RGCs induced at E15 but not P0 and adult is a result of genetic interactions between Ret and Brn3a, rather than a reflection of a normal change in expression patterns of Ret, Brn3a or both throughout development. Presumably, cell type specificity and/or Ret and Brn3a expression are determined by P0, such that after a loss of one or even both alleles of Brn3a in conjunction with the heterozygous state of Ret, the distribution of Ret^+^Brn3a^+^ RGC types is relatively unaffected.

### Sparse versus Complete double heterozygosity

In *Ret*^*CreERt2/WT*^; *Brn3a*^*CKOAP/WT*^ and *Ret*^*CreERt2/WT*^; *Brn3a*^*CKOAP/KO*^ mice, as sparse recombination is induced at either E15, P0 or adult, labelled RGCs lose one Brn3a allele in comparison to the surrounding tissue (Brn3a^AP/WT^ = “Het” RGC in a Brn3a^CKOAP/WT^ = “WT” territory, or Brn3a^AP/KO^ = “KO” RGC in a Brn3a^CKOAP/KO^ = “Het” territory, see Figure 1). If, as argued above, E15 induced Ret^CreERt2/WT^; Brn3a^AP/WT^ RGCs reveal a significant cell type distribution shift from the expected Ret^+^Brn3a^+^ fate, this would imply that the *Ret*^*KO/WT*^; *Brn3a*^*KO/WT*^ double heterozygote state is sufficient to alter developmental choices in RGCs. We therefore compared RGCs from *Ret*^*CreERt2/WT*^; *Brn3a*^*KO/WT*^; *Rosa26*^*iAP/WT*^ and *Ret*^*CreERt2/WT*^; *Brn3a*^*WT/WT*^; *Rosa26*^*iAP/WT*^ adult mice, in which recombination had been induced at E15. In this case, labelled RGCs are either double heterozygotes (Ret^CreERt2/WT^; Brn3a^KO/WT^) or single Ret heterozygotes (Ret^CreERt2/WT^; Brn3a^WT/WT^), respectively, and carry the same number of Ret and Brn3a alleles as the surrounding retinal tissue from conception into adulthood. Interestingly, we found that all RGC morphologies are consistent with the previously reported Ret expression domain, regardless of Brn3a dosage. Moreover, Ret^CreERt2/WT^; Brn3a^AP/KO^ RGCs labelled by P0 or P22 induction are effectively germline double heterozygotes before recombination, from conception to P0 or P22 respectively. These populations resemble the expected Ret^+^Brn3a^+^ expression domain (with the mentioned exceptions of ON and OFF β and ON Spiny RGCs), but are quite distinct from the E15 induced Ret^CreERt2/WT^; Brn3a^AP^ RGCs, regardless of Brn3a dosage. Combined, these observations strongly suggest that only early sparse but not complete *Ret*^*KO/WT*^; *Brn3a*^*KO/WT*^ combined heterozygosity can induce significant shifts in cell type distribution or morphological changes in RGCs. Potentially, Ret signaling in combination with information provided through Brn3a transcriptional control acts as a competitive factor in cell type specification of RGCs. Sparse double heterozygotes receive altered signals compared to the surrounding tissue, and therefore adopt novel cell fates or acquire altered abnormal morphologies. In the germline double or single heterozygotes, labelled RGCs and surrounding tissue have the same dosage of Ret and Brn3a, resulting in unaltered RGC type specification decisions. A similar phenomenon is observed in Purkinje neurons, which exhibit defects in dendrite morphogenesis in the presence of sparse but not germline knockout of the TrkC neurotrophin receptor ^59^.

### What are the molecular mechanisms of cross-talk between Ret and Brn3a?

In the somatosensory system there is a well-established role for neurotrophic signaling in neuronal specification, survival, differentiation, and neurite growth and branching ^24–27,38,56,60^. TrkA/B/C as well as Ret and its GFRα co-receptors are necessary for the specification of classes of nociceptors, mechanoreceptors and proprioceptors ^43–45^. Transcriptional regulation of neurotrophic receptors plays a major role in cell type specification of other projection sensory neurons ^61–63^. Brn3a and its family members are important regulators of development and specification of projection neurons in the Trigeminal Ganglion (TG), Spiral Ganglion (SG) and Dorsal Root Ganglion (DRG) ^19,22,56,64–70^, and their functions are mediated at least in part through regulation of neurotrophic receptors. In the *Brn3a*^*KO/KO*^ TG, TrkC expression is essentially lost from onset (E10.5), while TrkA and TrkB are initially (E10.5) expressed but turn off at E15.5, followed by extensive cell death in the TG. By E17, only 30 % of *Brn3a*^*KO/KO*^ TG neurons survive, a majority of which express the Ret receptor ^68^. These changes in neurotrophin receptors are accompanied by significant shifts in cell type distribution ^71–74^. In the spiral (acoustic) ganglion of *Brn3a*^*KO/KO*^ mice TrkC is downregulated resulting in dendritic arbor abnormalities ^67^. Brn3a loss of function also affects cell type distributions in a variety of DRG cell types ^75–78^, accompanied by dynamic changes in numbers of TrkA, TrkB and TrkC receptors, and increase in Ret^+^ cell numbers ^69^. Significantly, in DRGs, Brn3a is a direct transcriptional regulator of TrkA ^79^, while in RGCs Brn3a loss may modestly affect TrkA expression levels ^9,23^. In the retina, some TFs are shown to modulate neurotrophic signaling components. For instance, Dlx2, a known activator of Brn3b ^80^, also directly regulates TrkB receptor expression in RGCs ^81^. Conversely, instances of control of TFs by neurotrophins were documented in motor and sensory systems. Dendritic branching and connectivity of a subset of motor neurons in spinal cord is controlled by a TF encoded by the Pea3 gene, which is in turn induced by target derived GDNF signaling ^82^. In DRGs, GDNF activates a transcriptional program repressing neurite growth of sensory neurons ^83^.

Surprisingly, not much is known about Ret and Trk control of cell type specification and dendrite formation in RGCs. Gain and loss of function manipulations of the BDNF/NT4 – TrkB axis in mice and frogs did not result in RGC loss, however produced a range of phenotypes including changes in dendritic arbor formation and RGC axon shifts towards small diameter fibers^30,36,37^.

We now show that Ret, GFRα and Trk receptor expression levels are regulated in RGCs by Brn3 transcription factors (^9,23^ and Figures 7–9). In our hands, the numbers of Ret^CFP^, Brn3a and Brn3b-expressing RGCs are largely conserved between *Ret*^*KO/KO*^ (*Ret*^*CFP/CFP*^) and control animals. While our previous RNAseq experiments did not show a significant Brn3a-dependency of Ret gene expression in early postnatal age ^9,51^, we now find that in adults, Ret expression in the retina-specific Brn3a knockout is moderately but significantly altered. The partial redistribution of double positive (Ret^+^AP^+^) in controls to single (Ret^+^) cells in *Brn3a*^*KO/KO*^ retinas may be due to the loss of some (Brn3a^+^Ret^+^) - expressing neurons such as ON and OFF β and ON spiny RGCs. When compared to the wild type, *Brn3a*^*AP/KO*^ retinas exhibit a significant increase in GFRα1^+^, GFRα2^+^ and GFRα3^+^ cells in the GCL, at the expense of Brn3a^AP^ RGCs, potentially indicating that Brn3a expression suppresses Ret-GFRα expression in certain RGC types, leading to choices in cell type specification or morphological features in the dendritic arbors. On the contrary, numbers of TrkA^+^ Brn3a^AP^ and TrkC^+^ Brn3a^AP^ RGCs are somewhat reduced in *Brn3a*^*AP/KO*^ compared to *Brn3a*^*AP/WT*^ retinas, while TrkA^+^ and TrkC^+^ cell numbers in the GCL are increased. By analogy with the TGG and DRG systems, these shifts could induce alternative RGC cell type decisions.

Taken together, our data suggest that in the sparsely recombined Ret^CreERt2/WT^; Brn3a^AP/WT^ and Ret^CreERt2/WT^; Brn3a^AP/KO^ RGCs, the dosage reduction of Brn3a significantly affects the expression of signaling components of the Ret (Ret, GFRα-s or downstream molecules) and/or Trk pathway, that, when combined with the Ret heterozygosity, result in shifts in cell type specification or morphological defects. Ret can function as a competitive coreceptor for ligands involved in neuronal arbor formation and axon guidance, such as ephrin and p75-NTR ^84^ and Plexin /NCAMs ^85^, some of which are under transcriptional control of Brn3a ^9,23^. Thus, it is possible that the reduced Brn3a dosage in Ret^CreERt2/WT^; Brn3a^AP/WT^ and Ret^CreERt2/WT^; Brn3a^AP/KO^ RGCs results in partial loss of these co-receptors, and consequentially in morphological defects.

The proposed competitive nature of the Ret – Brn3a genetic interaction in the context of RGC specification could read out signals necessary to specify the appropriate numbers and spacing of distinct RGC types. Since both Ret and Brn3a are postmitotically expressed in RGCs, this could mean that RGC type fate is still plastic after exiting the cell cycle, as has been proposed for photoreceptors ^86–89^. This mechanism could then explain how individual retinal clones originating early in retinal development can adjust their composition to accommodate the diversity of RGC type distribution and density, by shifting cell type specificity according to local neurotrophic signaling originating from other already specified RGC types. Alternatively, target derived neurotrophic support could help eliminate excess RGCs, by engaging either Ret-GFRα or TrkA-C signaling in an activity or Brn3a dependent manner. Intriguingly, Ret ligands GDNF and Neurturin, and Trk ligands BDNF, NGF and NT3 are differentially expressed at relevant developmental stages in other retinal neurons, RGCs themselves or retinorecipient brain areas (Sajgo, 2017 and Supplementary figures 1–2). It will remain to explore which of these sources are relevant in the competitive mechanism we propose.

**Supplementary Table 1.** χ^2^ statistics for pair-wise comparisons between different sparse random recombination experimental groups (age and genotype) considering RGC subtype distribution

|  | P22 sparse Brn3a<br>HET vs KO<br>(Fig.3 b2) | P0 sparse Brn3a<br>HET vs KO<br>(Fig.3 b1) | E15 sparse Brn3a<br>HET vs KO<br>(Fig.3 b) | E15 complete<br>Brn3a HET vs WT<br>(Fig.6 b) | Sparse P22 vs P0<br>Brn3a HET<br>(Fig.3 b2 vs. b1) | Sparse P22 vs<br>E15 Brn3a HET<br>(Fig.3 b2 vs. b) |
| --- | --- | --- | --- | --- | --- | --- |
| ChiStat | 14.45 | 37.92 | 48.08 | 10.29 | 10.52 | 175.22 |
| P value | 0.071 | $3.9 \cdot 10^{-5}$ | $6 \cdot 10^{-7}$ | 0.328 | 0.231 | $1.6 \cdot 10^{-30}$ |
| Deg. of<br>freedom | 8 | 10 | 10 | 9 | 8 | 13 |
| Total<br>Nr. Cells<br>Cond1 | 148 | 89 | 324 | 169 | 148 | 148 |
| Total<br>Nr. Cells<br>Cond2 | 145 | 89 | 232 | 138 | 89 | 324 |

**Supplementary Figure 1.**
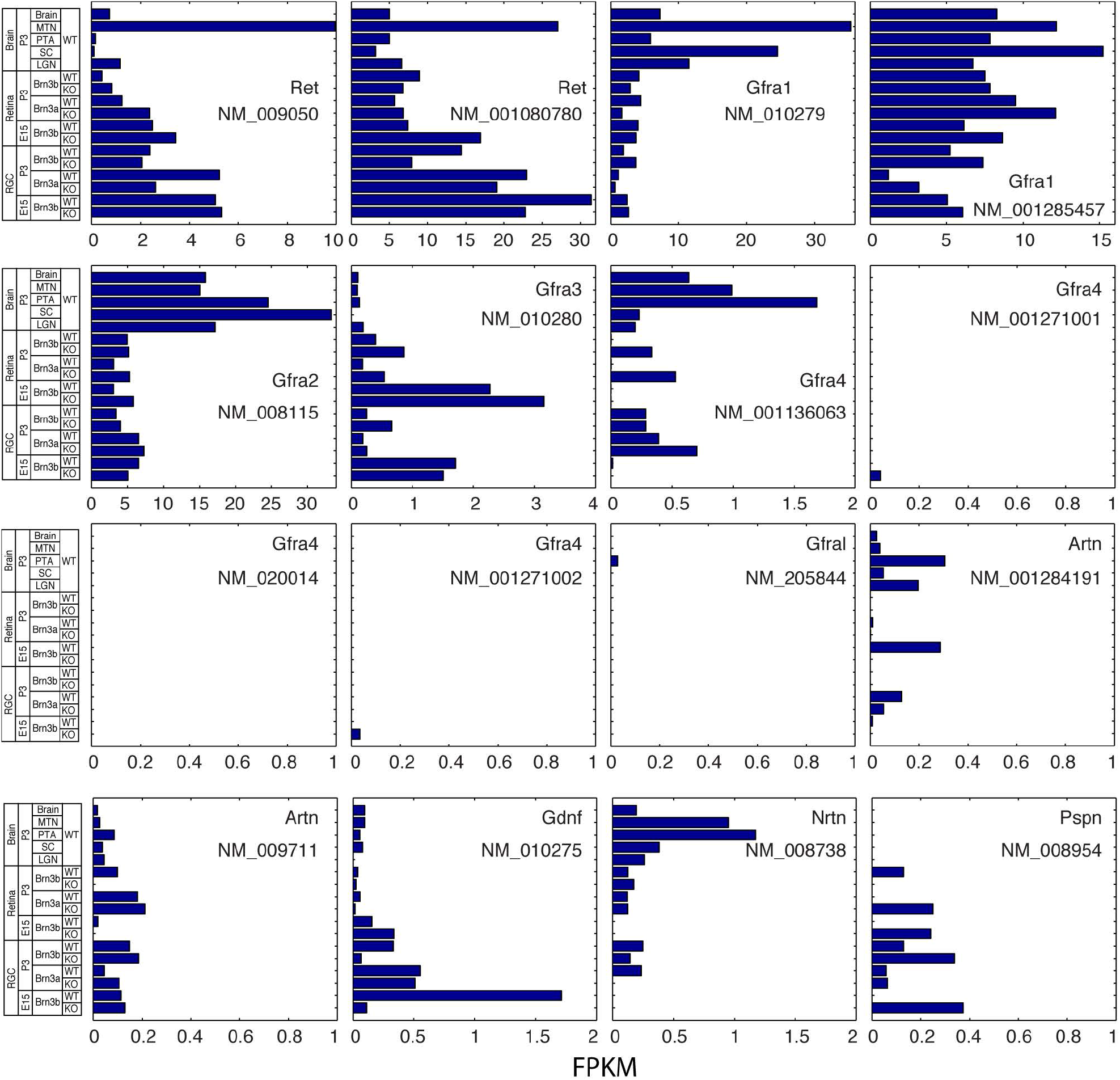
Expression of Glial Derived Neurotrophin Ligands and their receptors in E15 and P3 RGCs and major retinorecipient areas of the brain. Data are derived from Sajgo et al 2017. Each bar graph represents the FPKM (Fragments Per Kilobase of transcript per Million mapped reads) values for the indicated gene and RefSeq transcript (NM numbers indicated below the respective genes). Note that several genes have multiple transcripts, and only transcripts that were expressed in at least one of the samples are represented. Each bargraph scale is optimized to better reveal the expression levels of that particular sample. Brain samples are derived from WT P3 mice and represent from top to bottom, an average brain control sample, the Medial Terminal Nucleus (MTN), Pretectal Area (PTA), Superior Colliculus (SC) and the Lateral Geniculate Nucleus (LGN). Values for the brain areas represent medians for three samples (LGN: lateral geniculate nucleas, SC: superior colliculus), two samples (whole brain) or individual samples pooled from three animals (MTN: medial terminal nucleas and PTA: pretectal area). Retina and RGC samples are derived from either P3 or E15 mice, as indicated. Genotypes for retina and RGC samples are as follows: “Brn3bWT” = Pax6α:Cre; *Brn3b*^*CKOAP/WT*^; “Brn3bKO” = Pax6α:Cre; *Brn3b*^*CKOAP/KO*^; “Brn3aWT” = Pax6α:Cre; *Brn3a*^*CKOAP/WT*^; “Brn3aKO” = Pax6α:Cre; *Brn3a*^*CKOAP/KO*^. Retina values represent samples derived from two pooled retinas, while RGC values are medians of two biological replicates each derived from 6-8 retinas.

**Supplementary Figure 2.**
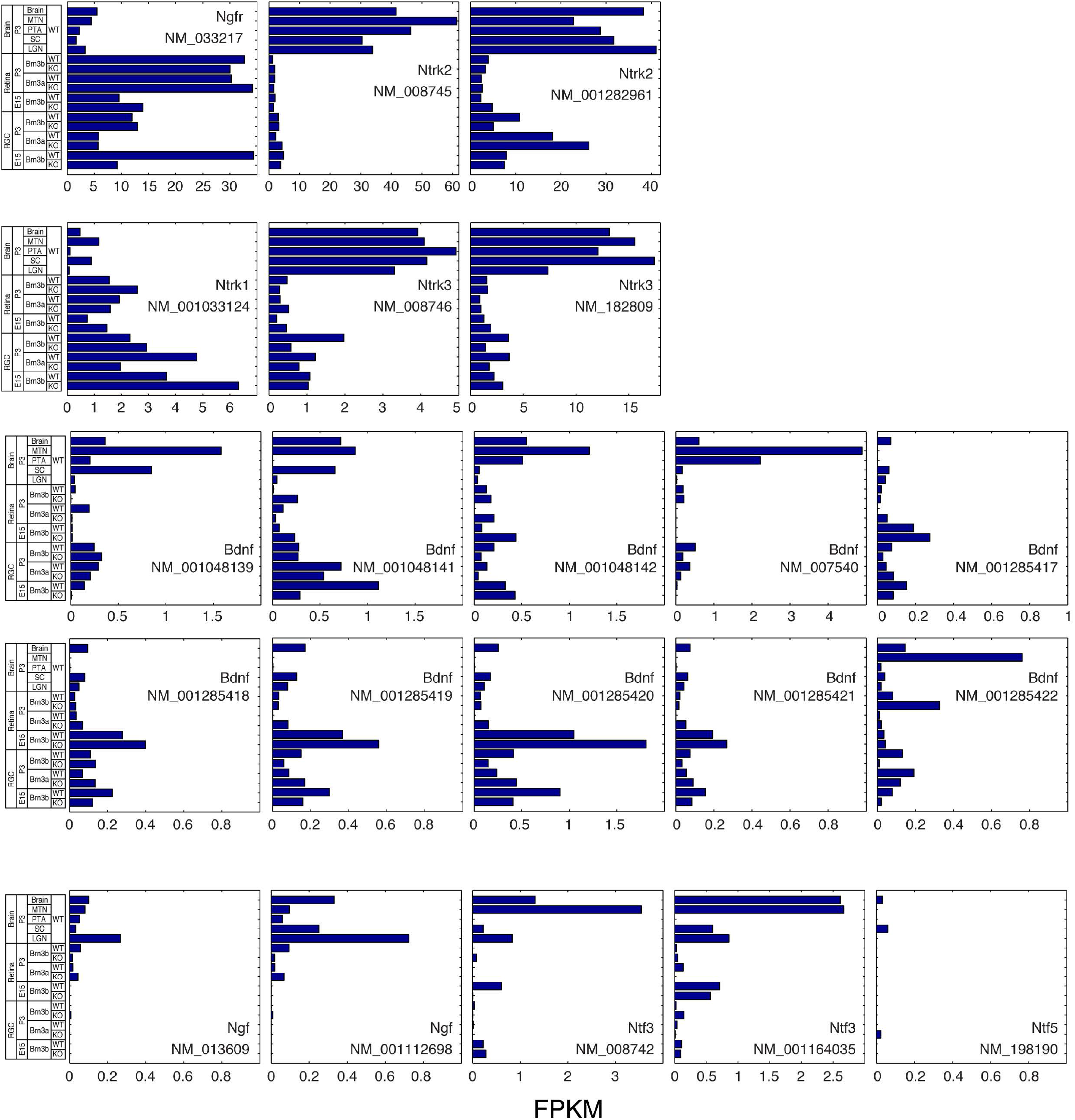
Expression of Target Derived Neurotrophin Ligands and their receptors in E15 and P3 RGCs and major retinorecipient areas of the brain. See Supplementary Figure 1 for data formatting, scaling and technical details.

## Acknowledgements

Wenqin Luo – U. Pennsylvania and Hideki Enomoto – Kobe University for helpful comments and providing mouse lines. Nadia Parmhans for assistance with Genotyping.

V.M. and T.B. designed experiments, prepared figures and wrote Manuscript. V.M. performed all experiments and collected all data.

The authors declare no competing interests.

All data related to this manuscript will be made available upon request from the corresponding authors.

